# Genotypic-phenotypic landscape computation based on first principle and deep learning

**DOI:** 10.1101/2023.02.09.527693

**Authors:** Yuexing Liu, Yao Luo, Xin Lu, Hao Gao, Ruikun He, Xin Zhang, Xuguang Zhang, Yixue Li

## Abstract

The relationship between genotype and fitness is fundamental to evolution, but quantitatively mapping genotypes to fitness has remained challenging. We propose the Phenotypic-Embedding theorem (P-E theorem) that bridges genotype-phenotype through an encoder-decoder deep learning framework. Inspired by this, we proposed a more general first principle for correlating genotype-phenotype, and the Phenotypic-Embedding theorem provides a computable basis for the application of first principle. As an application example of the P-E theorem, we developed the Co-attention based Transformer model to bridge Genotype and Fitness (CoT2G-F) model, a Transformer-based pre-train foundation model with downstream supervised fine-tuning (SFT) that can accurately simulate the neutral evolution of viruses and predict immune escape mutations. Accordingly, following the calculation path of the P-E theorem, we accurately obtained the basic reproduction number (*R*_*0*_) of SARS-CoV-2 from first principles, quantitatively linked immune escape to viral fitness, and plotted the genotype-fitness landscape. The theoretical system we established provides a general and interpretable method to construct genotype-phenotype landscapes, providing a new paradigm for studying theoretical and computational biology.

## Main

Due to the large amount of scientific data left to us, the SARS-CoV-2 virus has become a model virus for basic research on the transmission mechanism of virus-related epidemics, despite the Covid-19 pandemic having ended. To detect the evolutionary mutations that drive virus “immune escape”, researchers have employed various high-throughput experimental techniques [1,2]. However, these techniques have faced challenges in considering the possibility of epistasis and recombination across virus strains and in directly linking “immune escape” mutations to virus fitness. Now the GISAID database [https://gisaid.org] has collected more than 15 million SARS-CoV-2 genome sequences with spatio-temporal information. This massive data set provides an opportunity to uncover virus epidemic-related fitness information. Such findings can then be used to provide evidence for further experimental verification.

B. Hie et al. have made groundbreaking strides in the field of virus sequence analysis [3] by utilizing a natural language model. F. Obermeyer et al. [4] developed a hierarchical Bayesian multinomial logistic regression model, PyR0, to estimate the per-lineage fitness of SARS-CoV-2. M. C. Maher et al. [5] developed statistical models incorporating epidemiological covariates to account for the effects of driver mutations and the associated fitness of different viruses.

Our work focuses on constructing a computable representation of the genotype-fitness landscape by deciphering the effects of epistasis and recombination, identifying and predicting viral mutations associated with immune escape and quantifying virus fitness. To do this, following the fitness definition in population genetics, we first developed the P-E theorem, a theoretical framework that allows us theoretically to quantitatively calculate the fitness of a virus population by selecting appropriate embedding expressions for the virus genome under the Encoder-Decoder Seq2Seq framework. Then, on the basis of the P-E theorem, a more universal first principle of Genotype-Phenotype for landscape computing was proposed, and on the basis of the P-E theorem, we provide a computable basis for Genotype-Phenotype landscape computation under the deep learning framework. By calculating the transmissibility *R*_*0*_ of SARS-CoV-2, a macrobiological phenotype determined by the genotype of the virus, we give an example of constructing a real genotype-fitness landscape following first principles by using the P-E theorem. We then constructed a Transformer-based pre-train foundation model with downstream supervised fine-tuning (SFT) to calculate and predict virus immune escape mutations and the fitness without introducing and artificially coupling macro-epidemiological information, resulting in a precise mathematical representation of the virus genotype-fitness landscape in embedding space. In our work, the fitness, a macroscopic biological variable, can be accurately described mathematically as the expectation of a latent variable function according to a hidden state distribution.

Our model can also be used as a generative model to predict the likely occurrence of “immune escape” mutations. Our prospective and retrospective calculations (Figure 3B) have confirmed these results.

## Results

### Phenotype-Embedding (P-E) theorem, first principles and constructing genotype-fitness landscape

Unravelling the evolutionary causes and consequences of the genotype-fitness landscape is a fundamental yet challenging task. The features of the genotype-fitness landscape profoundly influence the evolutionary process and the predictability of evolution. However, the impact of the genotype-fitness landscape concept on evolutionary biology has been limited due to the lack of empirical information about the topography of natural genotype-fitness landscapes and the precise and quantified relationship between genomic variation and fitness [6]. To address this issue, we propose a novel approach combining basic statistical theory and deep learning models to computationally quantify the genotype-fitness landscape of the SARS-CoV-2 virus based on its genome sequence.

Starting from the definitions of evolutionary biology [6,7], we derive a formal expression for the fitness of a virus population, which is consistent with the mathematical expression given by F. Obermeyer et al. [4]. Under the Encoder-Decoder Seq2Seq framework that satisfies the universality theorem of neural networks [8], we introduce the Bayes theorem and combine it with Gaussian Mixed Model (*GMM*) expansion and Expectation-Maximum (*EM*) algorithm to deduce the P-E theorem under the condition of linear expansion according to the virus lineage (see Box 1 and supplemental note 1, formulas (1-11)): “An observable macrobiological phenotype can be calculated under the Encoder-Decoder framework if we can find a reasonable embedded representation of the related microscopic genotype”.

The P-E theorem provides a mathematical basis to establish a deep learning model to calculate virus fitness. This means that under an Encoder-Decoder framework with reasonable embedded representations, we can use only viral genomic sequence data to accurately compute viral population fitness as an observable macroscopic biological phenotype, and construct the genotype-fitness landscape of viral populations (Figure 1A).

**Figure 1.**
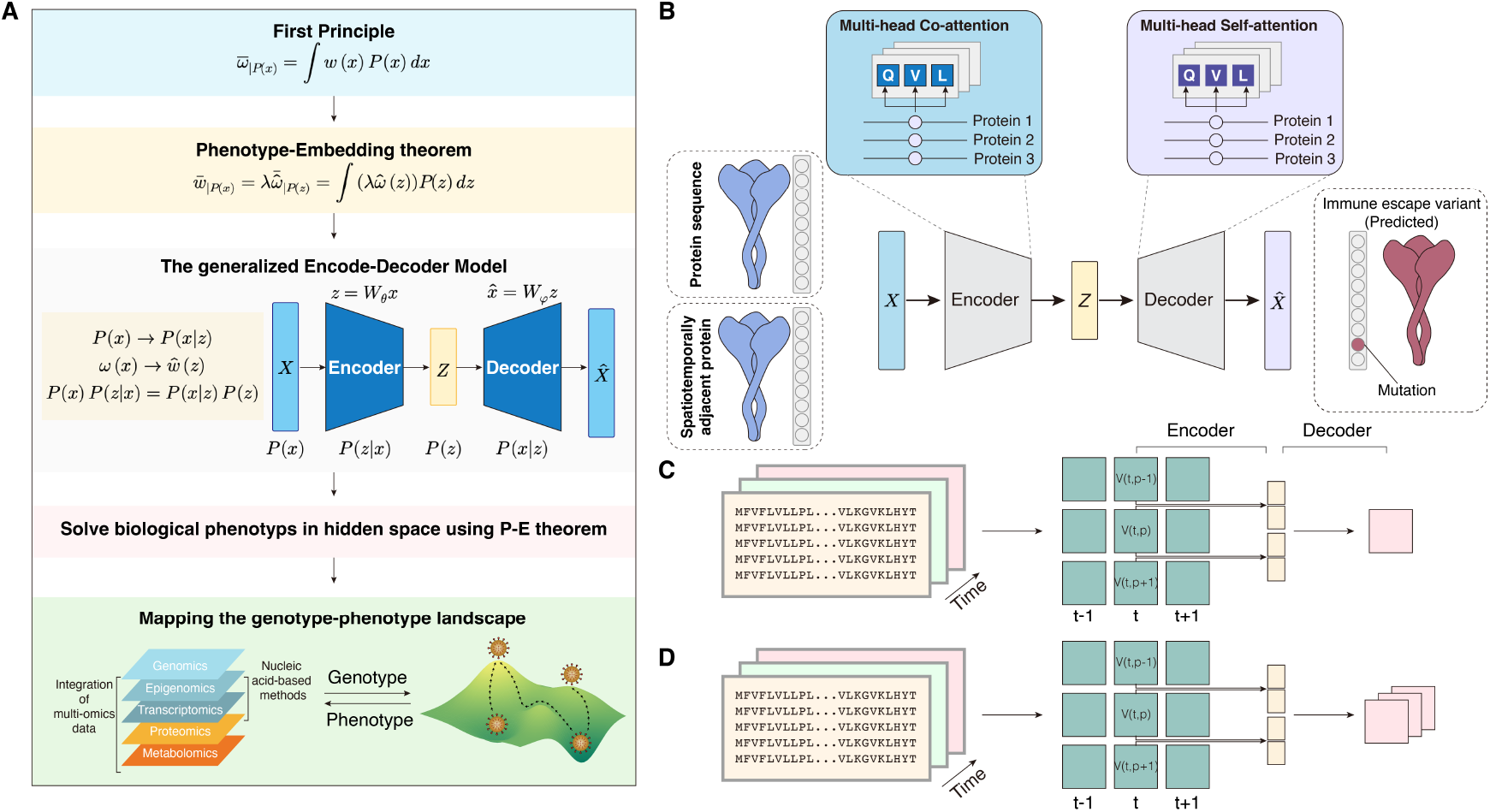
The P-E theorem, the framework of the CoT2G_F model and simulating the real evolution scenario of viruses. **(A)** Biological phenotype calculation processes. The process has five steps: 1. First principle, 2. The P-E theorem, 3. The generalized encoder-decoder deep learning model, 4. Solve biological phenotyps inhidden space using P-E theorem, 5. Mapping the genotype-phenotype landscape. **(B)** CoT2G_F is a Transformer-based pre-train foundation model with downstream supervised fine-tuning(SFT), it is a standard encoder-decoder architecture that introduces a co-attention and continuous span masking mechanism. **(C** and **D)** Shows the co-attention combined pre-training and fine-tuning steps of the model (see Supplementary Materials, Figure Extended Data 4).

Inspired by the P-E theorem and the formal logic of fitness calculation in the proof of the P-E theorem, we propose a more general first principle of genotype-phenotype: “A macroscopically observable biology variable can express as a mathematical expectation of a microscopic state function according to a state probability distribution” (see supplemental note 4, Formula (35) and (35)’). This “first principles” provides a mathematical basis for precisely defining biological phenotypes, allowing us to deeply understand and solve computational problems of biological phenotypes, and at the same time, the P-E theorem provides a computable basis for the application of first principles. On the basis of this work, we further deduce how to start from the first principles of universality and combine the P-E theorem to achieve a more general quantitative calculation framework of biological phenotypes (see Figure 1A). With the first principles and the P-E theorem, we can then design a suitable deep computational model to calculate the biological phenotypes of interest.

### Modelling of virus evolution mechanisms based on virus sequences

In recent years, natural language models have made great strides in learning the composition rules of DNA and amino acid sequences and exploring the evolution of biological species [3,9–11]. To effectively utilize the P-E theorem for fitness calculation, we need to devise a natural language model implemented within the Encoder-Decoder framework.

Different from B. Hie’s work [3], we construct a pre-trained + supervised fine-tuning(SFT) foundation model, a Co-attention-based Transformer pre-train model CoT2G-F to bridge Genotype and Fitness (see Figure 1): (i) Introducing co-attention and self-attention mechanisms [12] to extract the spatio-temporal correlation and long-range site interaction information within and across virus sequences. These mechanisms enable the model to extract epistatic and recombination signals that promote the occurrence of immune escape mutations. (ii) Dividing training process into two stages: pre-training, simulating the neutral evolution of the virus by randomly masking one or more continuous bases at any position of the input virus sequence, learning the “grammar” rules of sequence composition; and fine-tuning, in chronological order construct the evolutionary map of the virus sequences dynamically, perform supervised fine-tuning according to the spatial-temporal correlation constraints and target sequences, reproduce the role of environmental selection, bringing the model closer to the virus evolution process (Supplementary Figure 4).

Our approach stands out from other NLP models used in the study of viral sequence evolution primarily due to our optimized supervised fine-tuning (SFT) strategy. Specifically, we first start from the pre-training model, combine the P-E theorem to determine the mutation that contributes the most to the fitness of the virus as the target sequence, and then use the time-dependent dynamic fine-tuning mode to obtain the spatio-temporal selection signal of virus evolution that may not be directly attainable through other models (Figure 1C and 1D, Supplementary Figure 4A and 4B).

As of November 2022, 10.2 million SARS-CoV-2 spike proteins from GISAID (www.gisaid.org) were selected and subjected to strict quality control to train the CoT2G-F model (Extended Data Table 1). This data was used for pre-training, with further stages of fine-tuning to verify the model’s predictive ability. The CoT2G-F model can obtain the hidden state distribution of DNA bases or amino acids, and a sequence “embedding” that satisfies this hidden state distribution is guaranteed to conform to the “grammar” rules even in the presence of “semantic” changes. This allows for the identification and inference of “immune escape” mutations from the degree of “semantic” change (Figure 1B, 1C and 1B and see supplemental notes 2-4). Subsequent work will focus on using the CoT2G-F model to determine the micro-state function of a latent variable corresponding to the virus fitness in the hidden space, starting from the P-E theorem (see supplemental note 1, Formula (11)).

### Identifying and predicting “immune escape” mutations

Following the CSCS decision rule for semantics and syntax changes proposed by B. Hie [3], we tested the performance of our model CoT2G-F against two related mainstream models from the framework design in three technical dimensions. Our model demonstrated good performance (see supplementary materials). As shown in Figure 2A, our model outperformed the BiLSTM and Vanilla Transformer models, and the ability to identify and predict immune escape mutations was improved. By considering the spatio-temporal dynamics of virus evolution during training processes, the model was able to identify existing and predict future “immune escape” mutations. We tested this using more than 2 million SARS-CoV-2 genomic sequence data from the United Kingdom as an example (Figure 2B, 2C and 2D, Supplementary Figure 4, Figure 5C and 5D).

**Figure 2.**
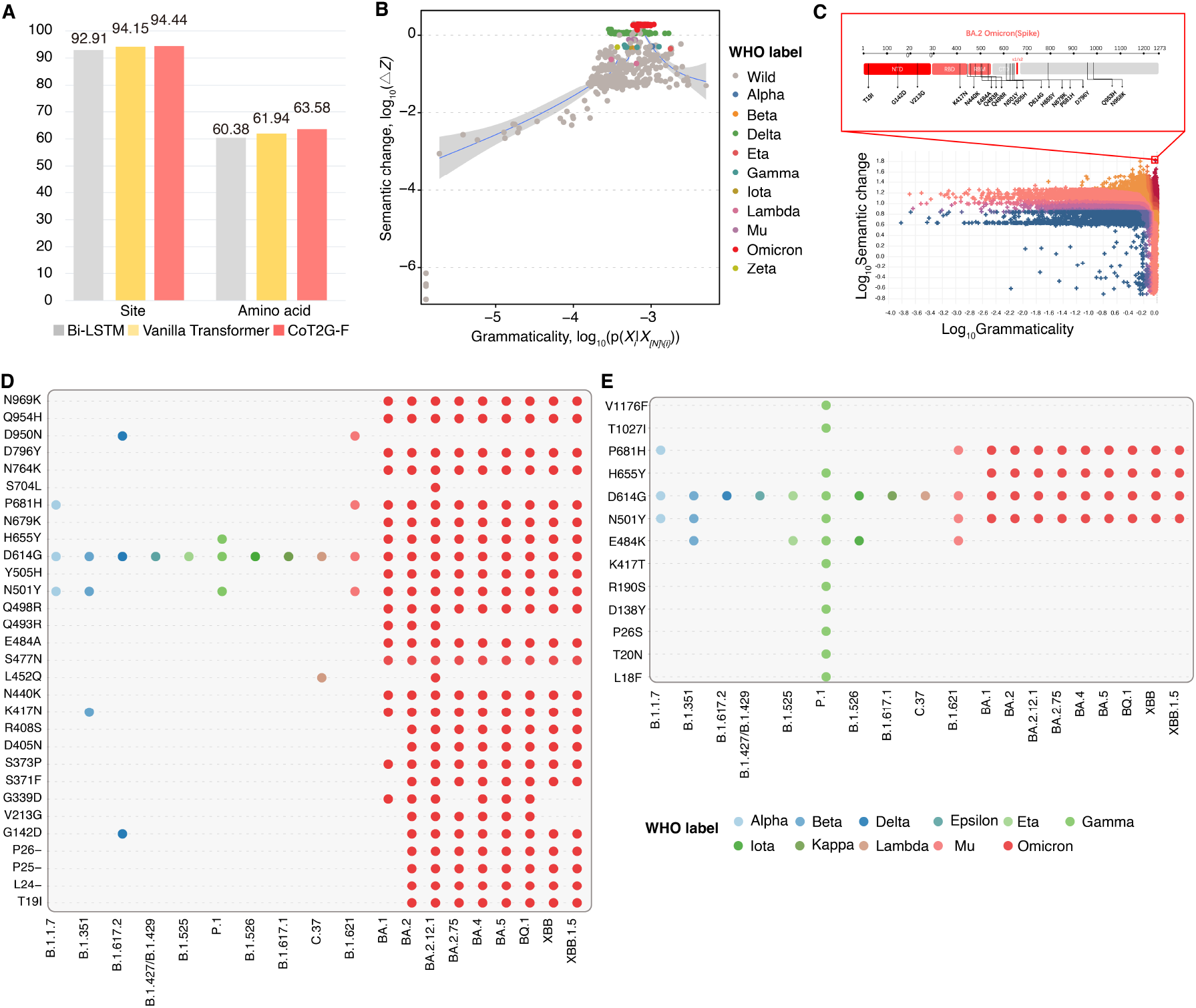
Modeling of virus evolution mechanisms based on virus sequences. **(A)** The figure shows a comparative experiment result among our method (CoT2G-F), Vanilla Transformer and Bi-LSTM (see supplementary materials). (**B**) The virus lineage with a high prevalence rate has a significantly higher semantic change score (CSC) (see supplemental note 4), calculated by the UK-submitted spike protein sequence data for June 2020 to February 2022. The left side of the figure (**C**) visualizes the semantic changes and grammaticality output by our model CoT2G-F with the horizontal axis representing the grammaticality and the vertical axis representing the semantic changes. The bottom side of the figure (C) visualizes the semantic changes and grammaticality output by our model CoT2G-F with the horizontal axis representing the grammaticality and the vertical axis representing the semantic changes. The upper right corner of scatter plot (C) means that the bigger the semantic changes and the more likely the virus strain is to immune escape. The upper side of the figure (C) shows the predicted mutation sites on the Spike protein sequence of the Omicron B.A.2 virus lineage with high semantic changes and grammatical fitness, these mutation sites are real and marked at the bottom of the right figure. (**D**) Take the virus sequence data collected in the United Kingdom from August to October 2022 as input to identify and reproduce emergent “immune escape” mutations. (**E**) Input sequences collected from the United Kingdom between March 2022 and June 2022 were selected to predict immune escape mutations.

Convergent evolution is a widespread phenomenon observed throughout the evolutionary process, including the later stages of viral epidemics. It refers to the independent emergence of similar traits or adaptations in different virus lineages, often in response to similar environmental pressures or functional requirements. This evolution typically occurs under strong environmental selection constraints [13]. Results showed that our model can predict and reproduce the convergent evolutionary mutations. For instance, nine of the 18 convergent mutation sites in the representative lineages of pre-Omicron and Omicron [13–15] were predicted with the model (Supplementary Figure 6).

Notably, our model was fine-tuning trained using data from GISAID as of March 31, 2022, demonstrating its ability to give accurate predictions up to six months in advance without dynamic fine-tuning. Besides, our model can also be used as a generative model to generate possible immune escape mutations for screening and early warning study (Supplementary Figure 6).

### Deciphering the intrinsic correlation between “semantic” change-related “immune escape” mutations and virus fitness

The immune escape ability of a virus lineage determines its fitness or the relative basic reproduction number (*R*_0_). The problem is establishing a quantitative relationship between the “immune escape” mutation and the fitness of the virus conforming to the P-E theorem under the framework of the natural language model. If evolution by natural selection is to occur, there must be a genotypic change in the virus population across generations. This change dependent on the genotype; if the genotype does not change, there is no change in fitness. Therefore, “semantics” and “grammar” together determine the virus’s fitness. Under the theoretical framework of deep learning, the above property can be expressed mathematically as the convolution of “semantic” and “grammatical” terms of the CSCS criterion [3]. When multiplied by the coefficient *λ*, the final mathematical representation is consistent with the P-E theorem (see Box 1, Formula (9) and Supplemental Note 1 and Note 4, Formulas (11) and (31)). Based on the above reasoning, we can argue that F. Obermeyer et al. [4] defined virus per-lineage fitness as precisely a special case of the P-E theorem. These results suggest that the term related to the “semantics” change of the model CoT2G-F and the CSCS criterion [3] may be the micro-state function of the latent variable corresponding to the virus’s fitness that we hope to obtain.

### Viewing the correlation between the trends of virus epidemics and the absolute mean distribution of conditional semantic change score *CSC*

The P-E theorem proves that we can construct the function of a latent variable through the Encoder-Decoder Seq2Seq model or its extension model. The mathematical expectation of this function in terms of the hidden state distribution is the corresponding macroscopic observable variable that we hope to obtain. We can extend and define a Conditional Semantic Change score (*CSC* score) from the *CSCS* score presented by B. Hie et al. [3]. The Conditional Semantic Change score (*CSC*), as a function of latent variables describing “semantic” changes, is used to measure the “immune escape” ability related to virus sequence mutation. Thinking that the absolute mean of the *CSC* score is an index of the immune escape ability of a virus lineage, we want to see how this index changes across different virus lineages (see Supplemental Notes 2-4, Formula (27)). We selected 2.7 million SARS-CoV-2 spike protein sequence data from the UK to calculate the absolute mean distribution of the *CSC* score according to the hidden state probability distribution (Figure 3).

**Figure 3.**
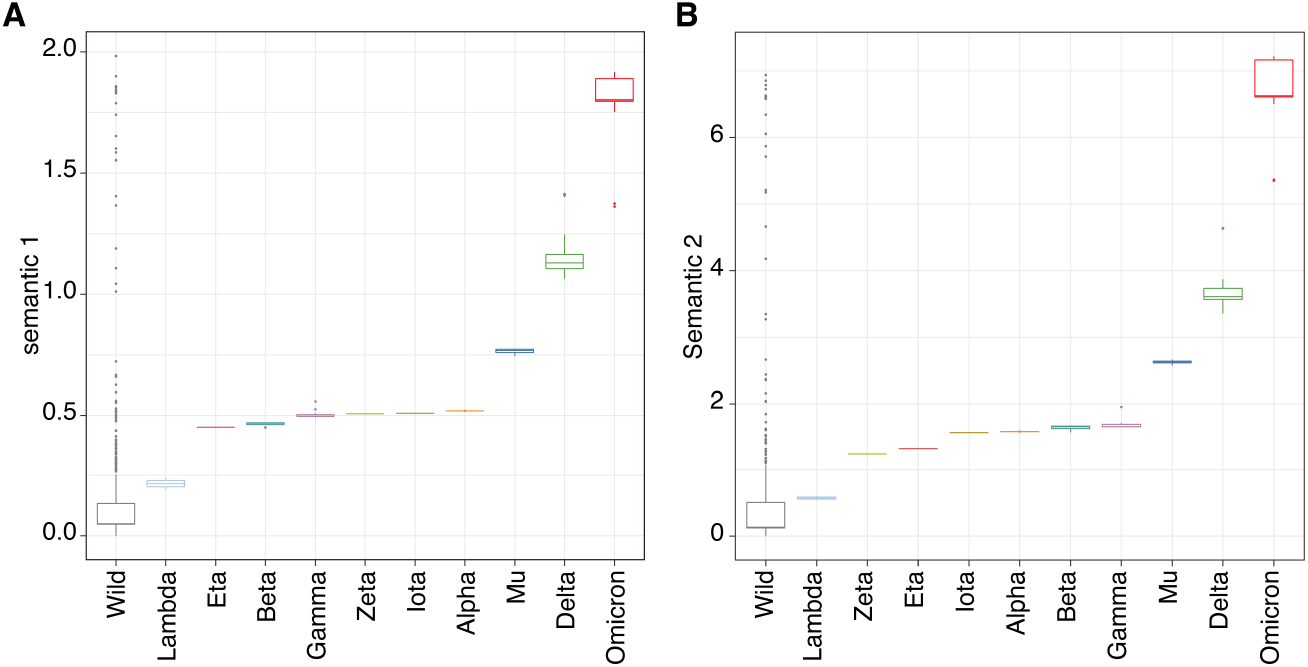
Absolute mean distribution plot of the semantic change parameter 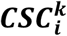. Take the virus sequence data collected in the United Kingdom after March 2022. (A) Using the *𝓁*_1_ norm. (B) Using the *𝓁*_2_ norm. The horizontal axis in the figures is the different virus lineages according to the emerging chronological order. The vertical axis is the absolute mean of the semantic change 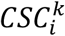 of the *k*_*th*_ virus lineage, which is defined as the immune escape capacity of the *k*_*th*_ virus lineage. Figures (A) and (B) both clearly show that with the progress of the global epidemic, from the Wuhan virus lineage to the Delta lineage and then to the Omicron lineage, the immune evasion ability continues to increase and even accelerates. Figures (A) and (B) show the same absolute mean distribution trend of the 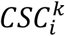, but figure (B) changes is more smoothly.

Figure 3A and 3B clearly show that, from the Wuhan lineage to the Delta lineage and then to the Omicron lineage, the immune evasion ability of the emerging virus lineages continues to increase and even presents an accelerated trend. The “immune escape” ability of the virus lineage is a calculable and quantifiable parameter directly related to the virus genome sequence mutation, which can be used to assess the cumulative collective effects of the immune escape mutations in an emerging virus lineage. The absolute mean of the *CSC* score accurately describes the changing trend of the “immune escape” ability of virus lineages and thus reveals the epidemic potency of the virus. This result further suggests that the *CSC* score is the microstate function corresponding to the virus fitness indicated by the P-E theorem.

### Expressing the fitness of a virus lineage as a convolution of “semantics” and “grammar” and the mathematical expectation of a semantic change function according to a hidden state probability distribution

It has long been a goal in biology to create genotype-fitness landscapes by mapping DNA sequences to mutation combinations observed in phylogeny or evolutionary experiments. However, due to the consideration of the epistatic effects of mutations, when the mutation number is large, the model becomes complicated and difficult to calculate and verify experimentally [6,7,16]. E. D. Vaishnav et al. [17] argue that the complete fitness landscape defined by a fitness function maps every sequence (including mutations) in the sequence space to its associated fitness. However, no theoretical model has yet been able to provide a concise computational model and framework for the genotype-fitness landscape while fully considering epistasis and recombination potential. Under the framework of the P-E theorem, we attempt to solve this problem through our deep learning natural language model CoT2G-F combined with *CSC* scores.

Based on the functional relationship between the *CSC* score and the latent variable of the CoT2G-F model (see Supplemental Note 4, Formula (22)), we know that the *CSC* score is a relative measure of semantic change corresponding to the relative immune escape ability of the virus and the “semantic” mutation corresponding to genomic mutation. Therefore, the *CSC* score measures the degree of the genomic mutation of emerging virus strains relative to wild-type virus strains. Furthermore, the *CSC* score and the “grammatical” coincidence degree determined by the mutation state distribution – the hidden state distribution – together form a measure of the virus’s immune escape ability in the form of convolution (see Supplemental Note 4, Formula (31)). Referring to the P-E theorem and by the properties of the *CSC* score and the formal equivalence of formulas (11) and (31), the *CSC* score should be the corresponding hidden space microstate function of the macroscopic observable variable *R*_0_. Formula (30) is the concrete realization of the P-E theorem (see formula (11)) under the CoT2G-F model. Finally, we acquire the basic formula for calculating *R*_0_ (see Supplemental Note 4 and Formula (33)). The correctness of the derived calculation formula for the per-lineage fitness is evident and verified (Figure 3A and 3B, Figure 4). This result and the P-E theorem provide a mathematical basis for redefining macro-phenotypes such as *R*_0_ and the resulting computability.

**Figure 4.**
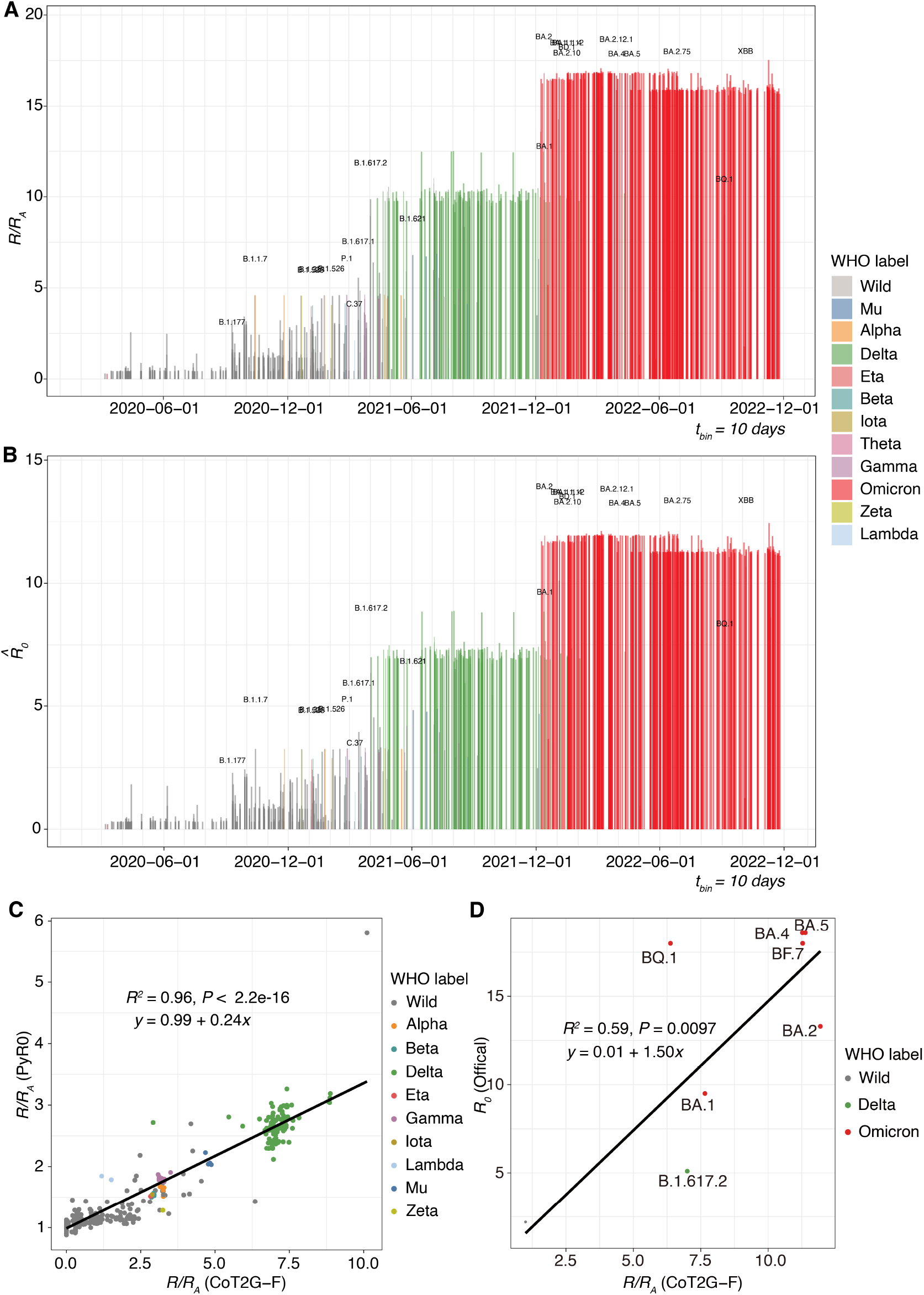
Relative fitness derived from the natural language model CoT2G-F versus date of lineage emergence. (A) This figure uses a time interval of 10 days. Referring to the Pango lineage designation and assignment, the relative basic reproduction number (*R*_*0*_), which is the fold increase in relative fitness of the virus lineages according to the Wuhan lineage, is plotted in different colours. The results for 5, and 20-day time intervals have shown in (Supplementary Figure 6). All the results are almost consistent, reflecting the robustness of the model. (B) The transmissibility potential (ie, absolute basic reproduction number 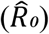 of virus lineages of SARS-CoV-2 by using an observable actual number of the absolute basic reproduction number 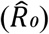 for the Wuhan lineage of SARS-CoV-2. (C) Comparison of the *R/R*_*A*_ calculated by the Co2TG-F with the values calculated with PyR0. (D) Comparison of the *R/R*_*A*_ calculated by the Co2TG-F with the official results.

### The genotype-fitness landscape

The calculation formula of the *R*_*0*_ has a novel meaning (see Supplemental Note 4, Formulae (31), (32) and (33)): it is the convolution of the semantic change function and the syntactic state distribution function in the embedding space. Here, convolution means giving the average of the cumulative effects of the contribution of all sampled virus variants to the *R*_*0*_ according to the reference sequence (we selected Wuhan-Hu-1 as a reference sequence; see Supplementary Materials). More importantly, the *R*_*0*_ can be precisely rewritten as a mathematical expectation of a semantic change function according to a hidden state probability distribution, giving the quantitative relationship between the “immune escape” ability of the virus and the fitness of the virus lineage. Finally, we present a comprehensive and precise formulation of the virus genotype-fitness landscape. The consistency between these two different representations provides a solid mathematical basis and research paradigm for applying deep learning models to biological problems and gaining interpretability (see Supplemental Notes 1-4).

Studying the intrinsic relationship between statistical mechanics and deep learning has always been an intriguing subject [18–20]. We know that the core concept of statistical mechanics is: “The macroscopic physical observables can be characterized as the ensemble average of the corresponding microstate variable functions”. Inspired by this, the inference is naturally drawn: “Under the Encoder-Decoder framework, a macro-biological observable can be expressed as the mathematical expectation of a function of a latent variable according to a hidden state distribution in the decoder space”. We call this framework bridging genotype-phenotype landscapes is an important corollary of the P-E theorem and gives a novel interpretable application of deep learning theory to life science (Figure 5, Supplemental Note 1 and 4, equations (11), (35) and (36)). Finally, we can plot the genotype-fitness landscape in the embedded space based on the variation state of the viral genome sequence according to equation (30). It is a two-dimensional hypersurface, and we can obtain the fitness of the virus lineage by integrating the surface density of the specific region corresponding to the virus lineage on this hypersurface, thereby defining the immune escape force of the virus (see Supplemental Note 4, Figure 5B and Supplementary Figure 9).

**Figure 5.**
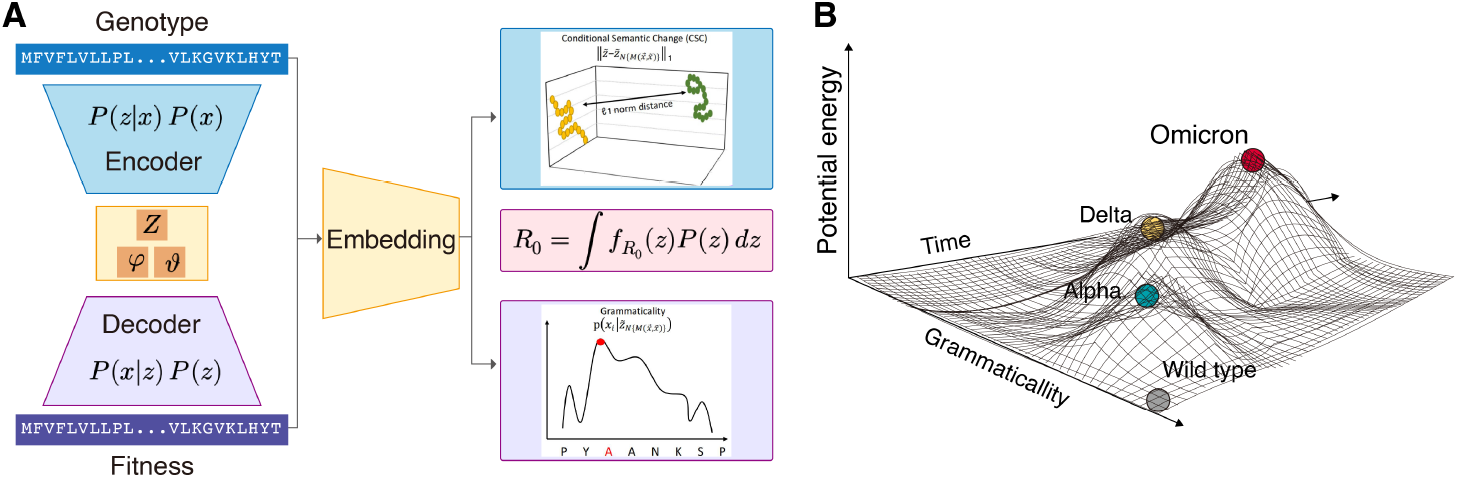
CoT2G-F framework for building genotype-phenotypic landscapes. **(A)** CoT2G-F is a natural language deep learning model that introduces co-attention and continuous span masking mechanism and takes the Transformer as a kernel. The model links the prior probability ***P(x)*** and hidden state distribution *P****(z***|***x)*** to the prior probability *P****(z***) and the posterior probability distribution *P****(x***|***z***) in the encoder-decoder space via Bayes’ theorem, mapping viral protein sequences as latent variables reflecting semantic and grammatical changes in the embedding space. Then refer to the semantic and grammatical changes to identify the virus sequence mutation. According to the hidden state posterior probability distribution *P****(z***)in the decoder space, the model can express the *R*_*0*_ as the mathematical expectation of a specific latent variables function related to viral sequence mutation according to a hidden state probability distribution, building a genotype-fitness landscape. (**B**) Plot virus genotype-fitness landscape. As a two-dimensional hypersurface, the genotype-fitness landscape can plot in a three-dimensional mapping space with “semantics”, “syntax”, and time as axes. Starting from the sampled data, we fitted a quadric surface function and drew a sketch of the Genotype-Fitness landscape, in which only three virus lineages, including Omicron, were shown.

### Inferring the *R*_*0*_ of the virus lineage

According to our theoretical framework, computing the fitness of each virus lineage and the fold increase in relative fitness to obtain the transmissibility potential,ie, basic reproduction number (*R*_*0*_) of SARS-CoV-2 requires adequate sampling for each virus lineage in a specific region and time interval. In this region and time interval, the virus begins to emerge, sustain its spread, increase in number, and eventually reach a plateau. Based on the biological definition of fitness, the selected sampling time interval needs to ensure the virus is transmitted for enough generations to reduce the violent fluctuation of the signal [4]. It is also crucial for determining the occurrence frequency of the dominant cluster of the different lineages and simultaneously calculating the absolute mean of the *CSC* score (see Supplemental Note 4, Formula (30), Figures S3A and S3B). Currently, the GISAID (www.gisaid.org) contains over 2 million genome sequences of SARS-CoV-2 from the United Kingdom [http://gisaid.org], and the spatio-temporal distribution of the virus lineages reflects the major trend of the world epidemic of SARS-CoV-2, which can serve as a model system for a reliable and feasible G-P principle.

We computed the virus per-lineage fitness using 2.7 million SARS-CoV-2 spike proteins from the GISAID database (www.gisaid.org) and adopted the Pango lineage designation and assignment. To calculate the factor *λ*_*k*_ in the per-lineage fitness (see Box 1, Formula (9) and Supplemental Note 4, Formula (30)), we first performed lineage assignment with Pangolin. We then selected the *k*_*th*_ lineage and its nearest sublineages to form a population, sampling for a given time interval *t*_*bin*_, and *λ*_*k*_ was the occurrence frequency of the *k*_*th*_ lineage in this small population. The calculation result is shown in Figure 4, where the time interval we chose was 10 days (see Supplementary Figure 6). Our model correctly inferred that the WHO classification variant Omicron lineages (Pango lineage BA.2.X) had very high relative fitness in a pandemic. It was approximately: ∼ 10-12 times higher than the original Wuhan and lambda lineages (Figure 4A, [95% confidence interval (CI) 16.78 to 16.84], Supplementary Figure 7), accurately predicting its rise in the spread regions.

In our study, we propose a simple calibration process to obtain the absolute basic reproduction number 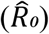 of virus lineages by using an observable actual number of the absolute basic reproduction number 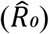 for the Delta lineage of SARS-CoV-2. (Figure 4A, Supplemental Note 4, Formula (34)). Our calculated absolute basic reproduction numbers 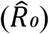 align with the results obtained by F. Obermeyer *et al*. [4] (Figure 4C) and are also in agreement with the actual and official monitoring results (Figure 4D). This result validates our proposed P-E theorem, allowing for the precise computation of the relative fitness of the virus lineage and *R*_*0*_.

## Discussion

Applying AI and deep learning methods to biology has two key points. The first is how to make the model more in line with biological logic, and the second is to delve into the connotation of the model and provide a mathematical basis for interpretability. This study explores these two aspects and presents an initial research paradigm. In the first step, we propose a general theorem, the P-E theorem, which is a computational path of macrobiological phenotypes under the framework of deep learning. Then in the second step, we constructed an encoder-decoder Seq2Seq deep learning model CoT2G-F that is more in line with the evolutionary biology scenario. The model can accurately predict the immune escape mutation of SARS-CoV-2 and calculate the immune escape ability of the virus lineage. In particular, because the CoT2G-F is a basic pre-trained foundation model for virus evolution research, the model can perform different downstream analysis tasks by various supervised fine-tuning (SFT).

In our work, The calculation of *R*_*0*_ was realized by the model CoT2G-F under the guidance of the P-E theorem, which gives the first principle-based interpretability of the model. The *R*_*0*_ is a mathematical expectation of the “semantic” change according to a hidden state distribution, which directly gives quantitative relationships among “semantics” change and “grammatical” fitness, “immune escape” mutation and the fitness of virus lineages (see Supplementary Materials, Supplementary Note 4, formulas (30) and (33)). This result can be seen as the correspondence in the biology of the core concept in statistical mechanics that “macroscopic observable physical quantities are the ensemble average of the corresponding microscopic state variable functions”, and is a corollary of the P-E theorem, which provides a new valuable perspective and interpretability for applying deep learning to virus evolution. It reveals the intrinsic correlation between deep learning theory and statistical mechanics studied by many researchers [18–20].

The practice and results of the *R*_*0*_ calculation in the present work provide a feasible paradigm for the computational modelling of genotype-phenotype landscapes and a compelling commentary on the first principle of “selection imprinting recorded in viral genomic mutation”, which is a concrete and successful example of the P-E theorem. We believe that, following the research path proposed in the present work, many research examples that adhere to the P-E theorem will emerge in the future.

In our study, we employ the Transformer as a kernel in our model, incorporating co-attention、self-attention and continuous span masking mechanisms to mimic the spatial-temporal dynamical associations of virus evolution. This comprehensive approach that combines the main deep learning algorithms provides several noteworthy benefits. Firstly, it enables us to extract long-range correlation mutation information from both upstream and downstream regions of the viral sequence, thereby capturing crucial spatio-temporal features relevant to virus evolution. Secondly, our model can effectively simulate genetic drift and discerns which mutations are more likely to be advantageous and subsequently fixed through natural selection, and exhibits significantly improved accuracy in the identification and prediction of potential future “immune escape” mutations. The efficacy of our model is clearly demonstrated in Figures 2A, 2B, 2C, and 2D, which provide compelling evidence of its predictive capabilities.

Importantly, we have constructed a formal three-dimensional embedding space to visualize the virus genotype-fitness landscape, utilizing the quantitative correlation between “immune escape” mutations and viral lineage fitness. This pioneering approach represents the first precise mathematical representation of a virus’ genotype-fitness landscape, enabling us to understand the topological properties that drive virus evolution and immune escape. The development of this fundamental methodology establishes a revolutionary theoretical and technical framework for future research, with potential applications across diverse contexts.

The proposed P-E theorem may demonstrate its universality in biological research by effectively studying fundamental phenotypic states in biology, such as brain homeostasis, metabolic homeostasis, cell fate state, ‘tumour microenvironment, survival capability, and advanced ageing state [21]. This theorem establishes a robust mathematical foundation for deterministically describing biological homeostasis and deciphering its stability mechanisms.

## Acknowledgements

We thank Guoping Zhao, Li Jin, Jiarui. Wu, Peiji Zeng, Ruibin Liu and Zhijian Lang for their deep and helpful discussions. We thank Tao Zeng and Junwei Liu for their assistance with the manuscript. We are also particularly grateful to Brian Hie for his constructive responses to our questions about the details of his related excellent models.

## Author contributions

Conceptualization: Yixue Li. Methodology: Yao. Luo, Yuexing Liu, Xin Lu, Hao Gao, Ruikun He, Xin. Zhang, Xuguan Zhang and Yixue. Li. Investigation: Yao Luo, Yuexing Liu, Xin Lu, Hao Gao and Ruikun He. Resources: Yixue Li. Data curation: Yuexing Liu, Yao. Luo, Xin Lu, Hao Gao and Ruikun He. Writing, original draft: Yixue Li, Yuexing Liu, Yao Luo, Xin Lu, Hao Gao. Writing, review and editing: Yixue Li, Yuexing Liu. Supervision: Yixue Li. Funding acquisition: Yixue Li.

## Methods

### Data access and preprocessing

Virial sequence and metadata were obtained from ViGTK (https://www.biosino.org/ViGTK), which is a mirror site for SARS-CoV-2 genome data of the GISAID (https://gisaid.org) and NCBI (https://www.ncbi.nlm.nih.gov) database. We download 16,395,521 genome sequences from ViGTK as of November 26, 2022. Records with missing time or location were discarded. After removing low quality sequences of length less than 29000 bp or with ambiguous bases more than 400, we obtained 14,959,290 genome sequences altogether (Extended Data Table 1). Lineage assignment was performed with Pangolin (version 4.1.2) [22] subsequently. To get the spike protein sequences, genome sequences were firstly aligned to SARS-CoV-2 reference genome (Wuhan-Hu-1, NC_045512.2) [23] using the MAFFT7 (version 7.505) [24]. Then, coding sequences were extracted from the alignment and translated into protein sequences. We only considered spike protein sequences whose sequence length were between 1265 and 1275 amino acid residues. Additionally, spike proteins with ambiguous amino acids were excluded, which yielded high quality dataset of 10,184, 273 sequences in total (Supplementary Figure 1). A total of 2,713,327 SARS-CoV-2 spike protein sequences that belong to UK was subset from processed dataset.

### Preparation of the dataset for pre-training and fine-tuning

After preprocessing, we bin time intervals into one-month segments, resulting in 35 time-bins (Supplementary Figure 4). In each month, sequences were arranged by their evolution distance relative to Wuhan-Hu-1 strain (NC_045512.2) of SARS-CoV-2 (Supplementary Figure 5). In order to find nearest neighborhood for the sequence in *t* time, all sequences in *t-1* month were aligned to this sequence. The spike protein sequence in *t-1* month with minimum edit distance relative to Wuhan-Hu-1 strain were selected as the nearest neighborhood.

### Co-attention based Transformer model bridging Genotype and Fitness

We built a pre-train foundation model CoT2G-F with downstream supervised fine-tuning(SFT). It is an Encoder-Decoder Seq2Seq architecture natural language model with a Transformer as the technical framework and introduced the co-attention and contiguous spans masking mechanisms. Pre-trained with 10.2 million SARS-CoV-2 Spike protein sequence data from GISAID, the data set with 2.7 million sequences from the UK region was selected to validate our model. Details have been described in the supplemental notes below.

### Comparison with other major natural language models

Figure2A shows a comparative experiment result with our method (CoT2G-F) and Vanilla Transformer without the introduction of the cross-attention mechanism, and Bi-LSTM, which is used in B. Hie [3]. to capture the semantic and syntactic variations of protein sequences. It is worth noting that B. Hie (4) and our method (CoT2G-F) solve different problems: B. Hie [3]. uses Bi-LSTM as an encoder to extract semantic variation and syntactic information and predict Immune escape, and our objective is to generate the protein sequence of immune escape in the next time slice, so we connected the decoder module in Bi-LSTM to generate protein sequence. In this figure, the comparison experiment is based on Bi-LSTM, Vanilla Transformer, and our method (CoT2G-F). We compare the effect of the model from tow dimensions, the first is the identification of sequence mutation sites, we hope that the model can accurately predict the mutated sites (i.e., high recall) without randomly predicting the mutated sites (i.e., high precise), so the f1 score combine with recall and precise is used to compare the identification of mutated sites by different models, denoted as ‘Site’ in this figure, compared with Bi-LSTM and Vanilla transformer, our method (CoT2G-F) improved f1 score of mutation sites by 1.53% and 0.29% respectively. The second dimensions is to identify the amino acid mutated into the site based on the identification of the mutation site in the first dimension, denoted as ‘Amino acid’ in this figure, we find the effect of Bi-LSTM is not ideal when dealing with the task of generating long protein sequences, and cannot accurately identify the mutated amino acids. However, the attention mechanism in Vanilla Transformer and our method (CoT2G-F) can capture the long-range dependencies of long protein sequences and can learn the mutation logic of amino acids relatively better. Compared with Vanilla Transformer, our method (CoT2G-F) improves the identification rate of mutated amino acids by nearly 1.64%, thanks to the information gain brought by the introduction of spatiotemporal information in our method (Figure 2A).

### Model training based on the fundamental principles of evolutionary biology

CoT2G-F is an Encoder-Decoder Seq2Seq architecture natural language pre-training model (see supplemental note 1) that introduces a co-attention and contiguous spans masking mechanisms and uses a Transformer as a basic framework. To make the model better reflect the real virus evolution scenario, we adopt a two-step training mode.

The first step is the pre-training process. First, from the outbreak of the virus to the present, the time cross-section is divided into months, and the global SARS-CoV-2 genome sequences in one month are arranged according to the evolutionary distance to form a two-dimensional genome sequence matrix ***X***_***t***_ **= (*x***_(*t*,1)_, …, ***x***_(*t,p*)_, …, ***x***_(*t,N*)_), the row of the submatrix ***x***_***t***,***p***_ represents the genome sequence of a virus around spatial position *p*, and *t* is the corresponding time of a certain time cross-section. During pre-training, the co-attention module of CoT2G-F starts from the first row of the matrix ***X***_***t***_, and sequentially takes three spatially connected Spike protein sequences ***x***(_*t,p*_,_−1_), ***x***(_*t,p*_), ***x***(_*t,p*_,_+1_)), get (***x***(_*t,p*_,_−1_), ***x***(_*t,p*_)) and(***x***(_*t,p*_), ***x***(_*t,p*_,_+1_)) pairwise co-attention signals, and then the Transformer module randomly masks any amino acid sites of the Spike protein with a proportion under 15%, which is completely different from [4]. We adopt such a pre-training process that can not only replicate the “neutral evolution” source of virus variation but also consider the long-range associations between sites in the viral genome sequence composition rules, as well as possible recombination effects. Therefore, our pre-training model that reflects the “semantics” and “grammar” of viral sequence composition is more reasonable. After completing the first step of training, we can already calculate the “semantic” changes of the Spike protein of each input virus genome sequence as defined in (see supplemental note 4) and select the target sequences according to the *CSCS* defined in [4] for the second step training.

The second step of training is the dynamic fine-tuning process. The occurrence of virus “immune escape” mutations are the result of selection. Pre-training can also indirectly sense the evolution of a virus over time. It is certain that if the evolution direction of the virus over time is highlighted in the training, it will be able to get a stronger “selection” signal and can more accurately identify and predict “immune escape” mutations. According to the time arrow, we take the three adjacent virus genome sequence matrices ***X***_***t* −1**_, ***X***_***t***_, ***X***_***t* +1**_, the five selected sequences in ***X***_***t* +1**_ with the largest *CSCS* scores used as the training targets, and then the three spatially correlated sequences (***x***_(*t,p*_,_−1)_, ***x***_(*t,p*)_, ***x***_(*t,p*_,_+1)_) are selected according to the same pre-training mode in ***X***_***t***_, and finally in ***X***_***t***−**1**_, select a sequence ***x***_(*t* −1,*p*)_ with the closest evolution distance to the sequence ***x***_(*t,p*)_ in ***X***_***t***_, when calculating the pairwise co-attention mechanism between (***x***_(*t,p*_,_−1)_, ***x***_(*t,p*)_, ***x***_(*t,p*_,_+1)_), while computing the co-attention mechanism between ***x***_(*t,p*_,_−1)_ and ***x***_(*t,p*)_. The sequence input of the dynamic fine-tuning process is sequentially expanded in Spatio-temporal order, from the outbreak to the present. Finally, we obtained a trained model CoT2G-F that can accurately reflect the “mutation + selection” mechanism during virus evolution, can predict and identify “immune escape” mutations, and then compute *R*_*0*_ (see supplemental note 4).

### “Immune escape” mutations prediction and *R*_*0*_ inference

As in [3], our model CoT2G-F can construct a semantic and grammatical “embedded” representation for given virus genome sequences and even has significant advantages to extract the upstream and downstream long-range associated variation information of the sequence itself and obtain the epistatic effects of “immune escape” mutations and recombination signals of virus strains. The model identifies and predicts virus “immune escape” mutational patterns by the learned semantic and grammatical rules. We further prove that the relative basic reproduction number *(R*_*0*_) or a fold increase in relative fitness of a virus lineage over the Wuhan lineage as a macro-observable variable is a mathematical expectation of a latent variable function according to a micro-hidden state probability distribution (see supplemental note 4).

When computing *R*_*0*_, we find this microscopic latent variable function is what we define as the conditional semantic change score (*CSC*). In our model, the *CSC* score counts the semantic change of the embedded virus sequence, reflecting the immune escape ability of the virus strain (see supplemental note 4). We applied CoT2G-F to all publicly available SARS-CoV-2 spike protein sequences following the Pango lineage designation and assignment to identify and predict “immune escape” mutations and calculate the per-lineage fitness of the virus without introducing additional epidemiological parameters and then obtain the *R*_*0*_ of the virus lineage. The introduced conditional semantic change score 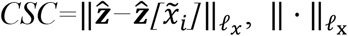 is the *𝓁*_*x*_ norm, *x* = 1, 2 (see supplemental note 1), in which 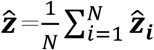. Considering the history of virus evolution and the reference of the relative fitness, we selected the virus sequence Wuhan-Hu-1 [GenBank: MN908947.3] as a reference corresponding to the latent variable 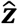 to align and normalize the CSC score (see supplemental note 4).

## Data Availability

GISAID sequence data is publicly available at https://gisaid.org. We used the following publicly available datasets for model training: SARS-CoV-2 spike protein sequences from GISAID (www.gisaid.org); training and validation datasets are deposited to https://zenodo.org/record/7388491. The data set sampled from the UK region according to the Pango lineage designation and assignment, used for the *R*_*0*_ calculation is available at https://zenodo.org/record/7388491.

## Code Availability

The source code for data preprocessing and modelling and available at https://github.com/martyLY/CoT2G-F.

## Key points

We present a precise formulation of the fundamental principles governing the calculation of biological phenotypes. Specifically, macroscopically observable biological variables (biological phenotypes) can be mathematically represented by the expectations of their corresponding microstate functions within their microstate distribution.

We introduce and substantiate the genotype-phenotype embedding theorem (P-E theorem). This theorem, established for the first time, demonstrates that the foundational principles of biological phenotype calculation can be accurately computed within the framework of deep learning.

Employing a deep learning model, we successfully calculate the fitness (*R*_*0*_) of SARS-CoV-2, an observable macro-phenotype. We provide a concrete example of biological phenotype computation.

The deep learning model we devised can effectively simulate the genetic drift and environmental selection involved in virus evolution. Furthermore, it accurately predicts the immune escape mutations of SARS-CoV-2.

## Tables

### Box 1

**The mathematical foundation and derivation of P-E theorem**

According to the basic definition of population genetics (*14*), the average fitness 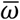 of a virus population can be expressed mathematically as follows:

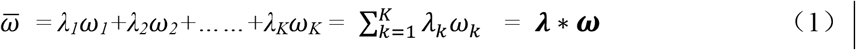

***λ*** = (*λ*_1_, *λ*_2_, … …, *λ*_*K*_), *λ*_*k*_ is the occurrence frequency of the *k*_*th*_ virus lineage in the whole population, which is an observable measure related to amino acid substitution characteristics, and ***ω***=(ω_1_, ω_2_, … …, ω_K_), ω_k_ is the fitness of the *k*_*th*_ virus lineage, and ***x =*** (*x*_*1*_, *x*_*2*_,…, *x*_*i*_,…, *x*_*L*_) represent L variation sites. Thus, each ω_k_ can involve the epistatic interaction of multiple mutation sites in the input sequence (*12*) as following:

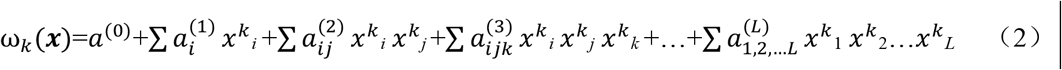

We define a function *P*(***x***) as a state probability distribution of the sequence’s variation states in real space. This variation causes changes in viral fitness. The *P*(***x***) can have an expanded form using a Gaussian Mixture Model (*GMM*).

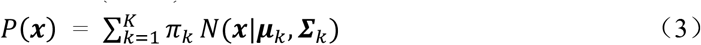

*π*_*k*_*N*(***x***|***µ***_*k*_, ***Σ***_*k*_) is the k_*th*_ component in the mixture model, *π*_*k*_ is the mixture coefficient, ***µ***_*k*_ and ***Σ***_*k*_ are the mean and variance values of the relative fitness, and they can obtain indirectly through the EM algorithm or deep neural network. Combining formulas (1) and (3), the formal representation of the average fitness of the virus population can obtain as follows:

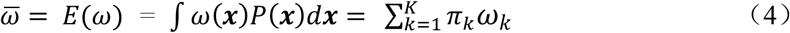

Let *λ*_*k*_ be the occurrence frequency of the dominant cluster in the *k*_*th*_ virus lineage when sampling virus strains, we can redefine the (dominant) per-lineage fitness as

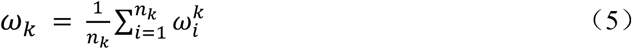

*ω*_*k*_ is a mathematical expectation of the contribution of a single virus strain to virus lineage or virus population (if *λ*_*k*_ = *π*_*k*_) fitness. Referring to formula (3) and (4),

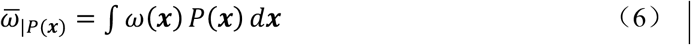

The subscript *P*(·) represents the state probability distribution corresponding to virus population. As long as we find a suitable functional form of *ω*(***x***) and a corresponding probability distribution *P*(***x***), we can obtain the per-lineage fitness of viruses by calculating their mathematical expectations by the formula (6), then get the average fitness of the virus population by formula (4). We can now try to do this within the context of deep learning, despite the fact that it is typically difficult to obtain and calculate *ω*(***x***) and *P*(***x***) directly from real-world data with noise. Considering the independent and identically distributed (*IID*) properties of state random variables of different samples in general, we let *P*(***z***|***x***) and *P*(***z***) be the hidden state probability distribution in encoder-decoder spaces under the Encoder-Decoder Seq2Seq framework, and *P*(***x***) and *P*(***x***|***z***) be the prior and posterior probability related to the input and output of the model. The relationship among *P*(***x***), *P*(***z***|***x***), *P*(***x***|***z***) and *P*(***z***) is given by Bayes’ theorem:

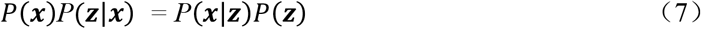

The function relationship defined by formula (1) still holds when *P*(***x***) *P* (***z***|***x***) maps to *P*(***x***|***z***)*P*(***z***). When the prior probability *P*(***x***) is unknown, obtaining the fitness of virus lineages in the encoder or decoder space is transformed into finding the hidden state probability distributions *P* (***z***|***x***) and *P*(***z***), the posterior probability distribution *P* (***x***|***z***) and the counterpart 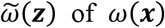.

Under the Encoder-Decoder Seq2Seq framework together with Bayes’ theorem and Universality Theorem of multilayer feedforward networks, because ***x*** *→* ***x****′* **=** *W****z*+***B*,

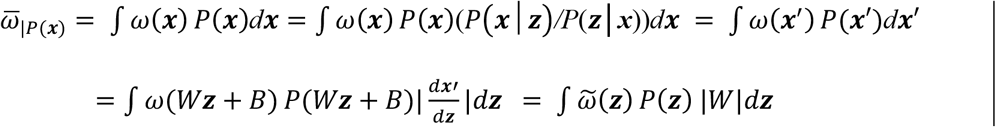

and 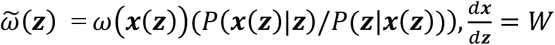 is the Jacobian matrix, and |*W*| is a determinant of Jacobian matrix that is the sum of combinations of products of elements in different positions of all the columns of matrix *W*. Considering the quasi-orthogonal property of base distribution in *GMM* expansion for ideal classification, let *λ*_*k*_ = |*W*|_*k*_ is a mapping of the |*W*| corresponding to the *k*_*th*_ virus lineages, and 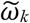 is a hidden space state function corresponding to the fitness of the *k*_*th*_ virus lineage. We set ***λ*** = (*λ*_1_, …, *λ*_*k*_) and 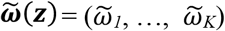 are vectors, then 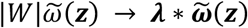. Finally, we have

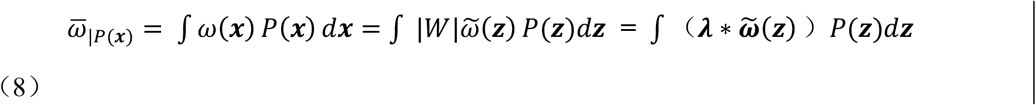

Obviously, formula (8) is the corresponding formal representation of formula (1) under the Encoder-Decoder Seq2Seq framework of deep learning, then we get a simple representation:

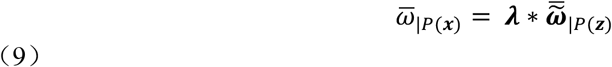

Because of the hierarchical structure of the population, lineage and cluster of the virus, and referring to the formula(1), (3), (4), (6) and (8), and based on the basic principles of population genetics (*14*), the elements of the vector ***λ*** are the corresponding elements of mixture coefficient in formula (4). The element *λ* is an occurrence frequency of a dominant cluster in virus lineage when sampling virus strains, and is a macroscopic parameter that links the hidden space with the actual space. Formula (9) gives a mathematical framework for representing phenotypes in embedded Spaces, leading to the Phenotype-Embedding (P-E) theorem: “An observable macro-biological phenotype can be computed under the Encoder-Decoder Seq2Seq framework if we can find a reasonable embedded representation of the related microscopic genotype.” Genotype-fitness landscape: we can plot the genotype-fitness landscape in the embedded space based on the variation state of the viral genome sequence. Starting from P-E theorem, we can regard the 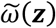 score as the “immune escape” potential of the virus, then, as a two-dimensional hypersurface, the genotype-fitness landscape can plot in a three-dimensional mapping space with 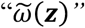, *“P*(***z***)”, and “time” as axes. The stable points on the hypersurface correspond to the genomic variation states that contribute the most to the “immune escape” ability. We can obtain the fitness of the virus lineage by integrating the surface density of the specific region corresponding to the virus lineage on this hypersurface. Now, if we take the gradient of the 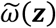 along the time and the *P*(***z***) axis on a two-dimensional hypersurface

## Supplementary Figures

**Supplementary Figure 1.**
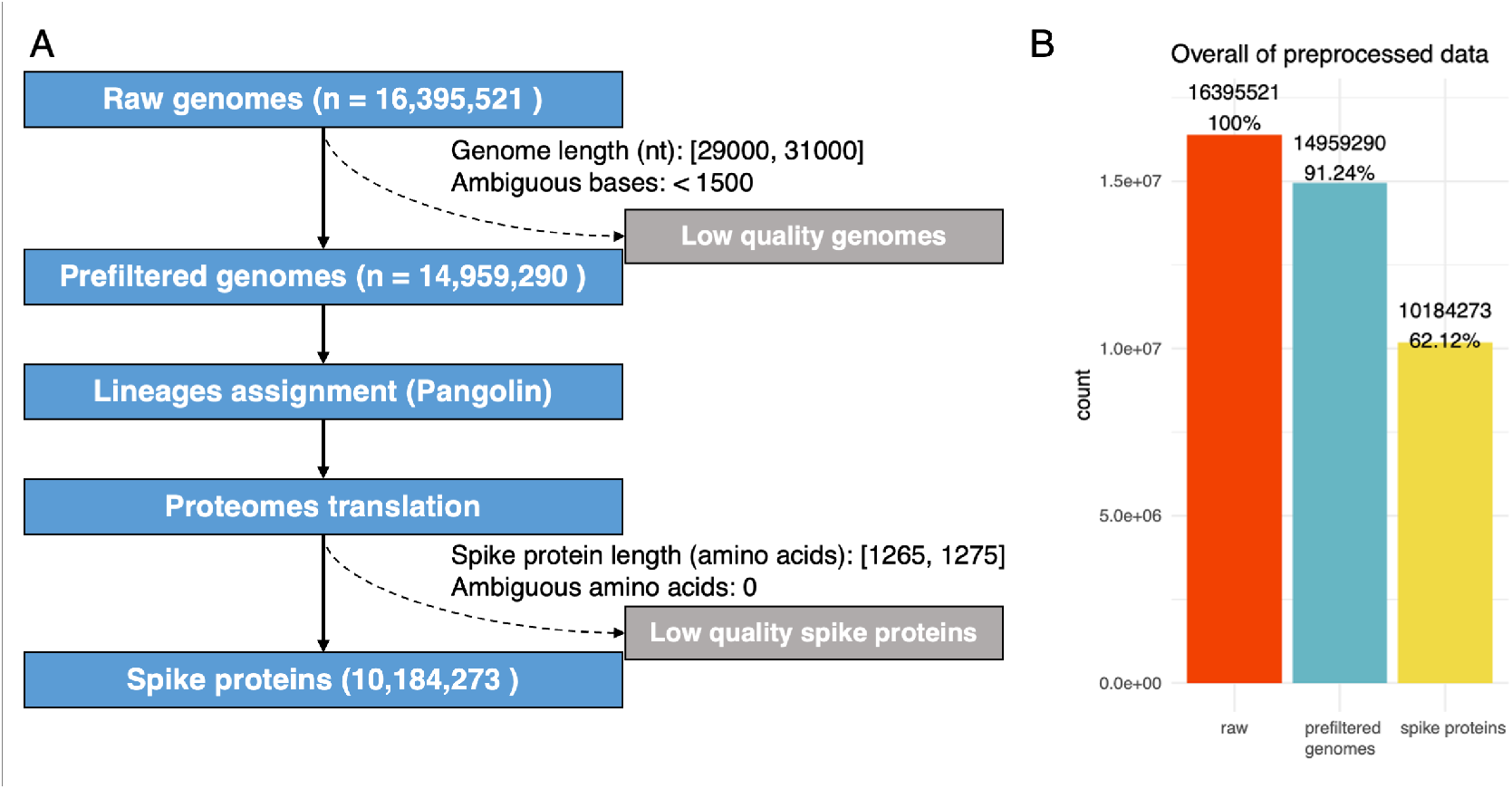
Preprocessing of SARS-CoV-2 sequences. **(A)** Workflow of the SARS-CoV-2 sequences preprocessing. **(B)** Summary of the preprocessed SARS-CoV-2 sequences.

**Supplementary Figure 2.**
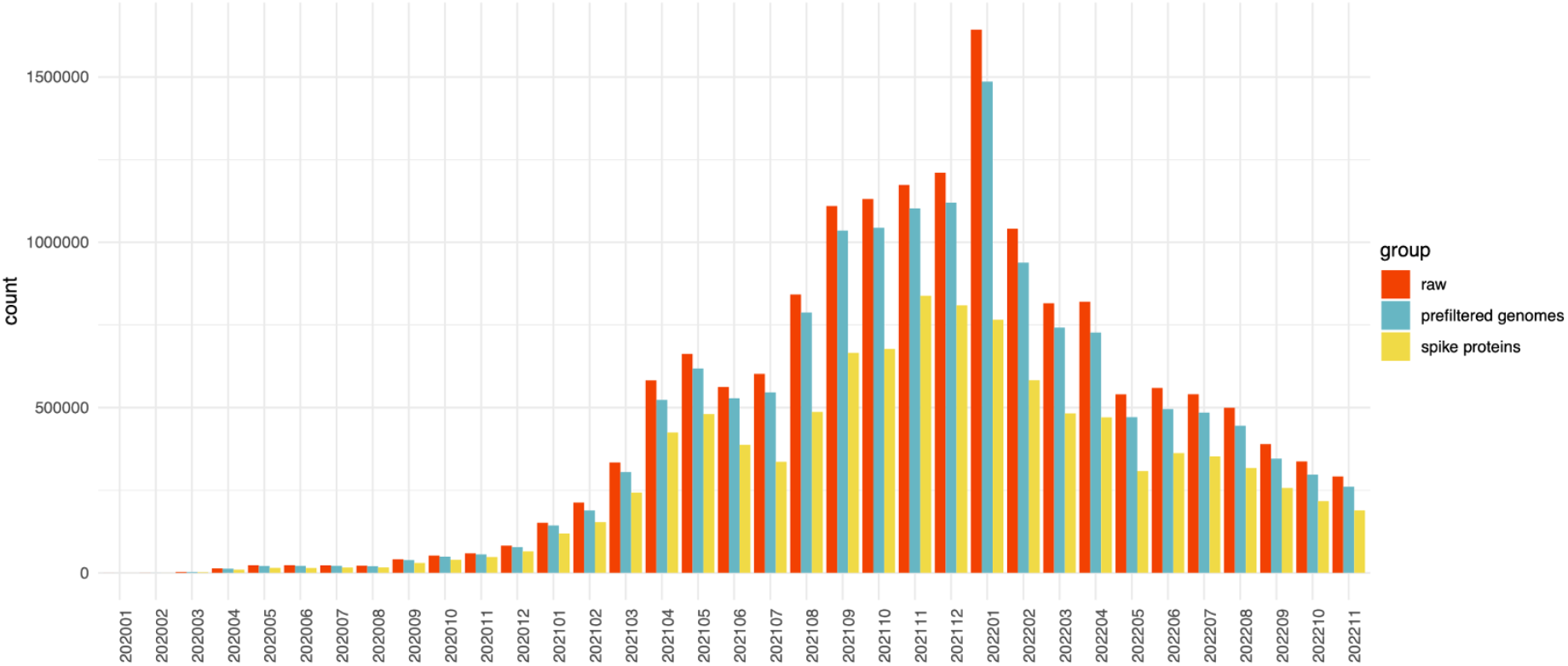
Summary of the preprocessed SARS-CoV-2 sequences.

**Supplementary Figure 3.**
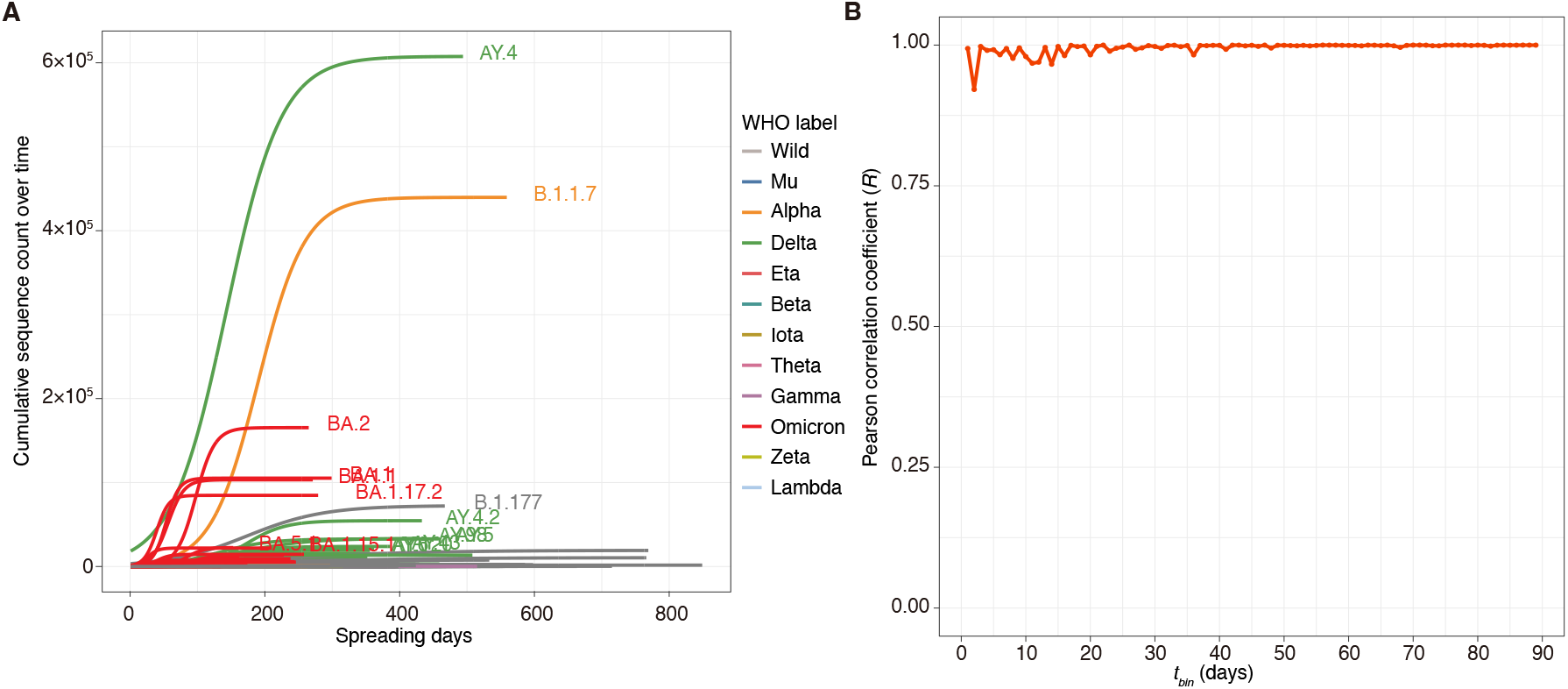
Growth rate of representative SARS-CoV-2 lineages in the UK. **(A)** Growth rates of SARS-CoV-2 lineages were fitted with a logistic model [1]. Only representative lineages were shown. From the above logical growth curve of each virus lineage relative to the spread time in the figure, a reasonable sampling time interval *t*_*bin*_ can be selected for calculating the occurrence frequency of the dominant cluster in different lineages. As can be seen from (A), starting from the first day to the time less than 100 days appears to be a reasonable time interval for sampling virus strains and calculating the occurrence frequency of the dominant cluster in different lineages. (B) The different t_bin_ versus the correlations among the distributions of the occurrence frequencies of the dominant cluster in different lineages. It can be seen that after 5 days the correlation is over 0.94, and around 50 days the correlation is close to one and keep stable. Based on (A) and the observed robustness of the correlations from (B) we can easily select any day between 5-50 days as the end point of the time interval *t*_*bin*_.

**Supplementary Figure 4.**
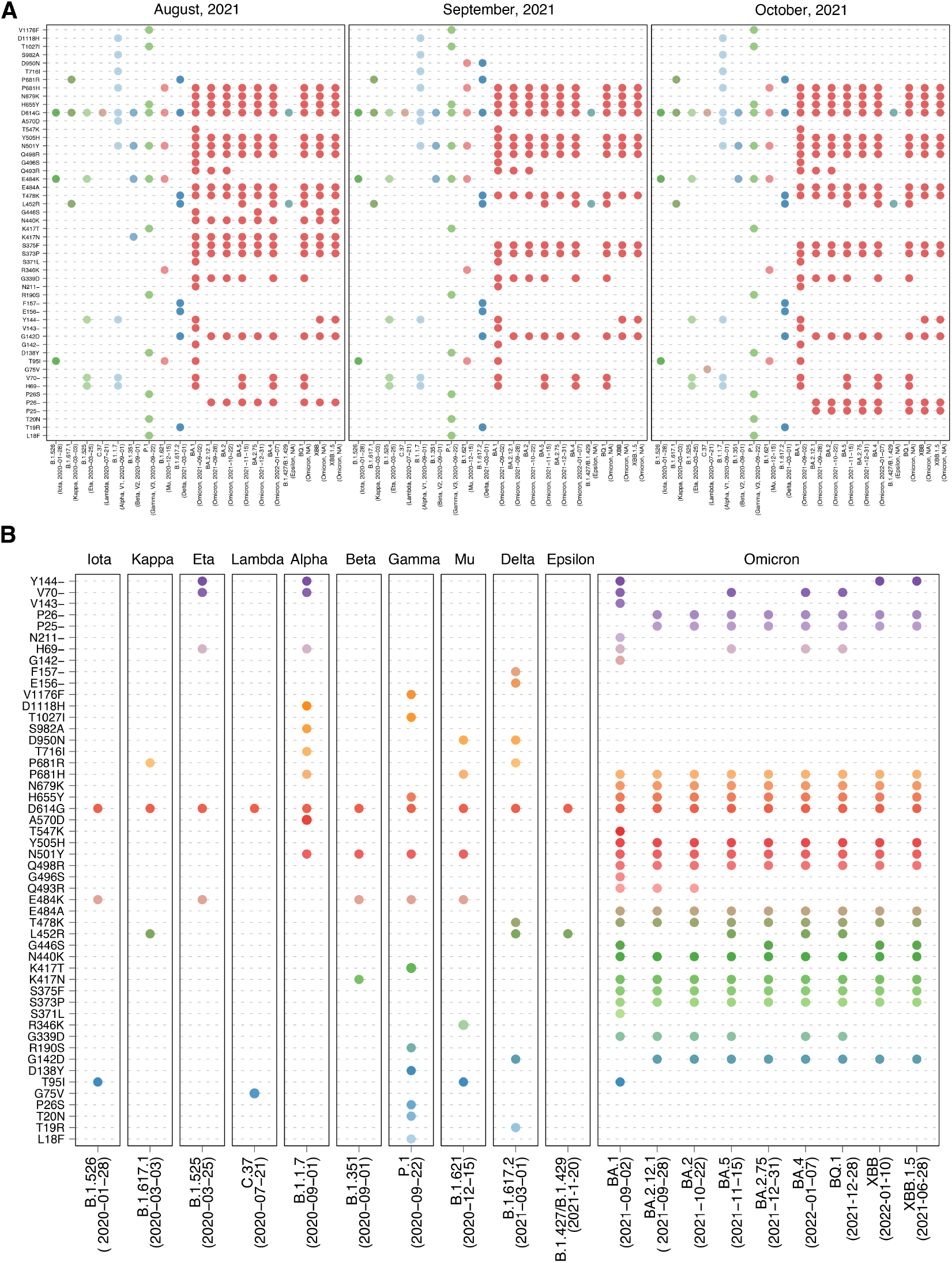
Reproducing known immune escape mutations. The prediction results show that the model has good reproducibility for already emerged “immune escape” mutations. We took data from August 2021 to October 2021 in GISAID (www.gisaid.org). The data were collected and submitted to the database by the United Kingdom.

**Supplementary Figure 5.**
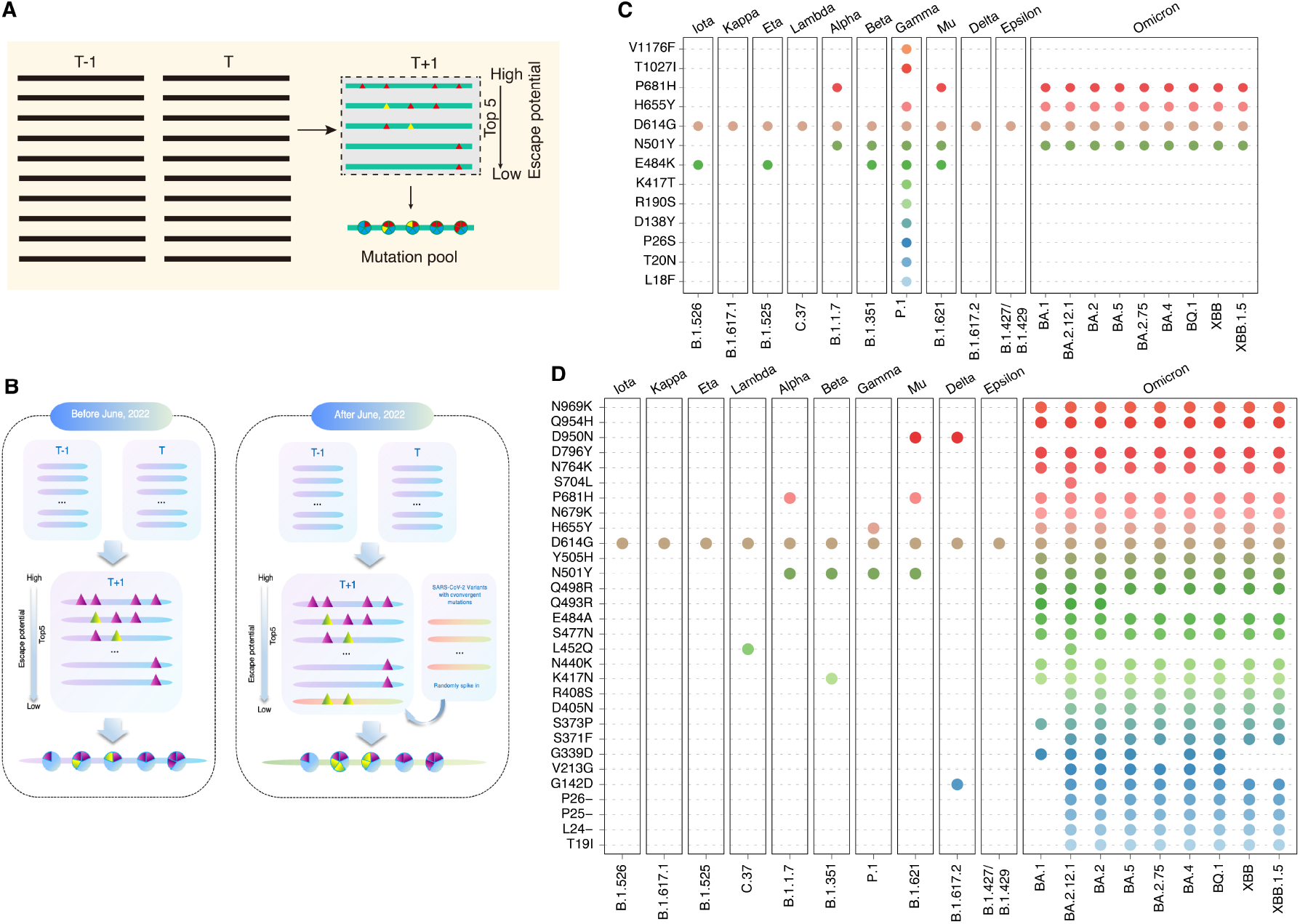
The schematic diagram for the target sequences search of the dynamic fine-tuning step. (**A**) The model selects virus sequences from three continuous cross-sections time, T-1, T, and T+1 time for the dynamic fine-tuning. Find the five virus sequences at T+1 time whose mutation sites are the key mutations that determine the virus lineage and whose sequences best meet the semantic and grammatical criteria. The five sequences are selected targets for one of the input sequence, and then supervised fine-tuning begins with the input virus sequence at T-1 and T time. In this fine-tuning process, at the same time, we have calculated the Spatiotemporal correlation among virus strains either. (**B**) Fine-tuning of targeted hot spot mutation areas was conducted from March to June 2022. Randomly added sequences containing convergent evolutionary sites were incorporated into the pre-training model. From June to November 2022, input sequences from the UK were selected for predicting immune escape mutations. (**C**) Prediction of immune escape mutations using data submitted to GISAID from the United Kingdom between March and June 2022. (**D**) The immune escape mutations including the convergent evolutionary mutations were reproduced.

**Supplementary Figure 6.**
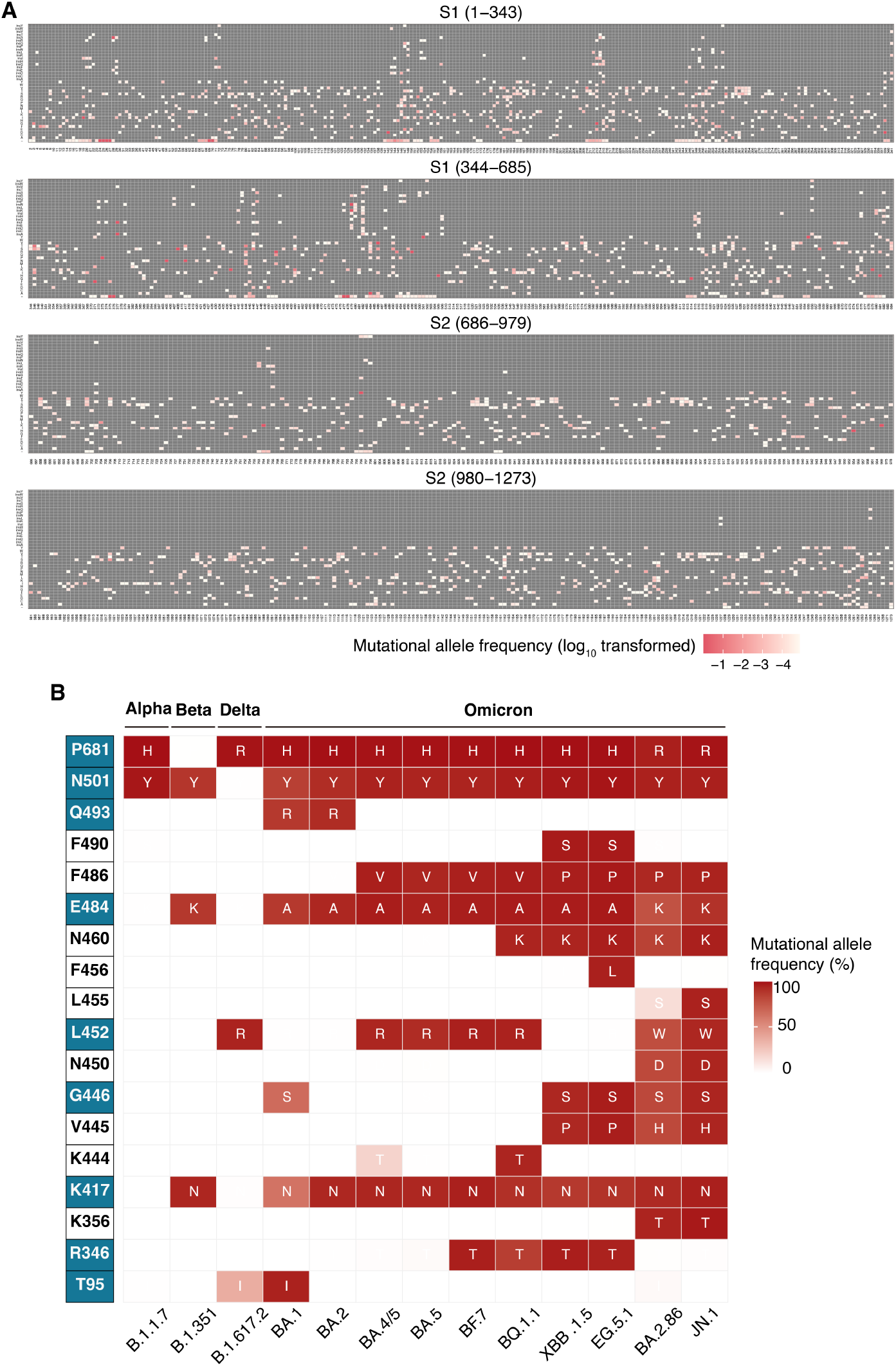
Model-Generated Spectrum of Immune Escape Mutants of SARS-CoV-2 and convergent evolutionary mutations. (**A**) The predicted immune escape mutant spectrum of SARS-CoV-2, including the three variants of the recently emerged virus lineage XBB.1.16. Heatmaps illustrating the predicted mutations across the Spike protein. The vertical coordinate stands for the possible mutations at each site, including amino acid substitute, insertions and deletions; while the horizontal coordinate represents for all positions (from 1 to 1273) in Spike protein. Squares are colored by mutational allele frequency (log_10_ transformed) according to the scale bar on the right. The model uses data from Britain from June to November. (**B**). The convergent evolutionary mutations predicted with the CoT2G-F. The heatmap illustrated the convergent mutations of the representative lineages of pre-Omicron and Omicron. The convergent mutation sites predicted with the CoT2G-F were highlighted in blue on the left side.

**Supplementary Figure 7.**
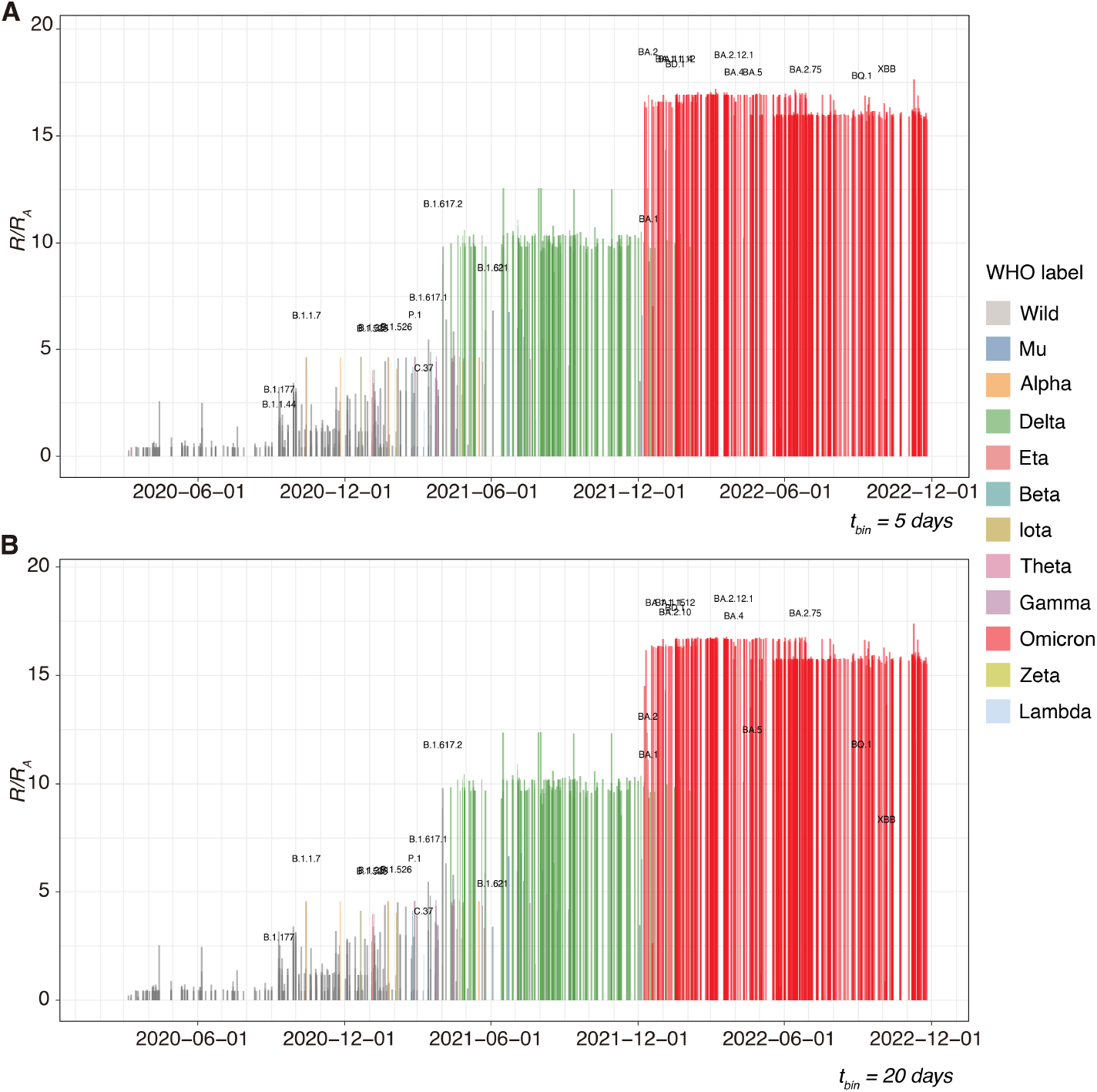
Relative fitness derived from the natural language model CoT2G-F versus date of lineage emergence. Circle size is proportional to the sampling number for different lineage in certain time intervals (*t*_*bin*_), which reflects the prevalence of the virus. Using the WHO classification, the *R*_*0*_, ie. the fold increase in relative fitness of the *k*_*th*_ virus lineage according to the Wuhan lineage is plotted in different colours. Figures A and B use 5-day and 10-day sampling time intervals to calculate *R*_*0*_, respectively. The results from A and B are almost completely consistent, reflecting the robustness of the model.

**Supplementary Figure 8.**
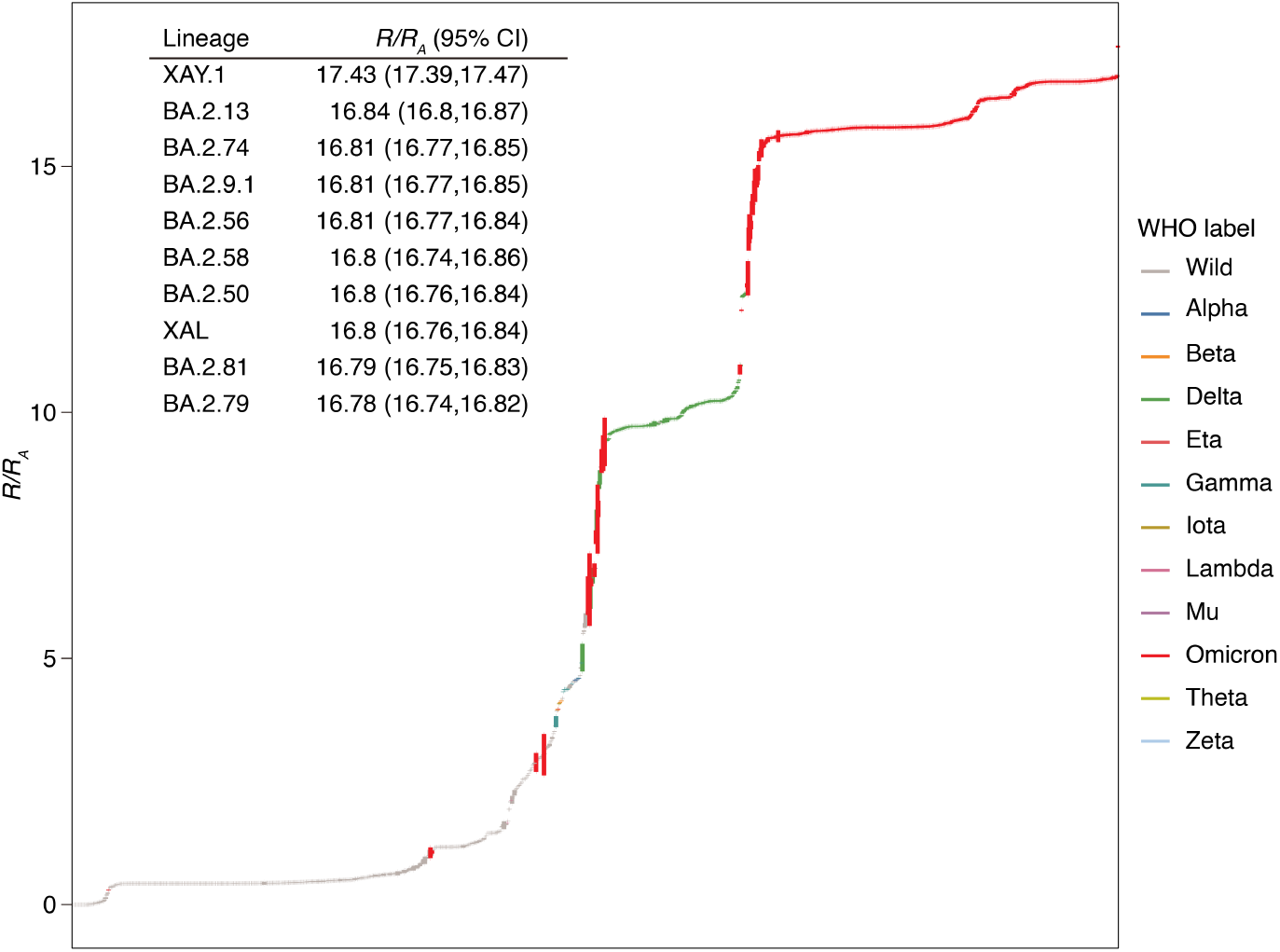
The confidence interval (CI) distribution for the *R*_*0*_ calculation. We select the sampling time interval (*t*_*bin*_) from 1 day to 60 days to calculate the R_0_ of the virus lineages, and the curves give the 60-day average of the *R*_*0*_. This curve also reflects the robustness of the model to calculate *R*_*0*_. The inset table lists the confidence interval (CI) of the ten fittest lineages of the pandemic inferred by our model. *R/R*_*A*)_ is the fold increase in relative fitness over the Wuhan lineage in the UK.

**Supplementary Figure 9.**
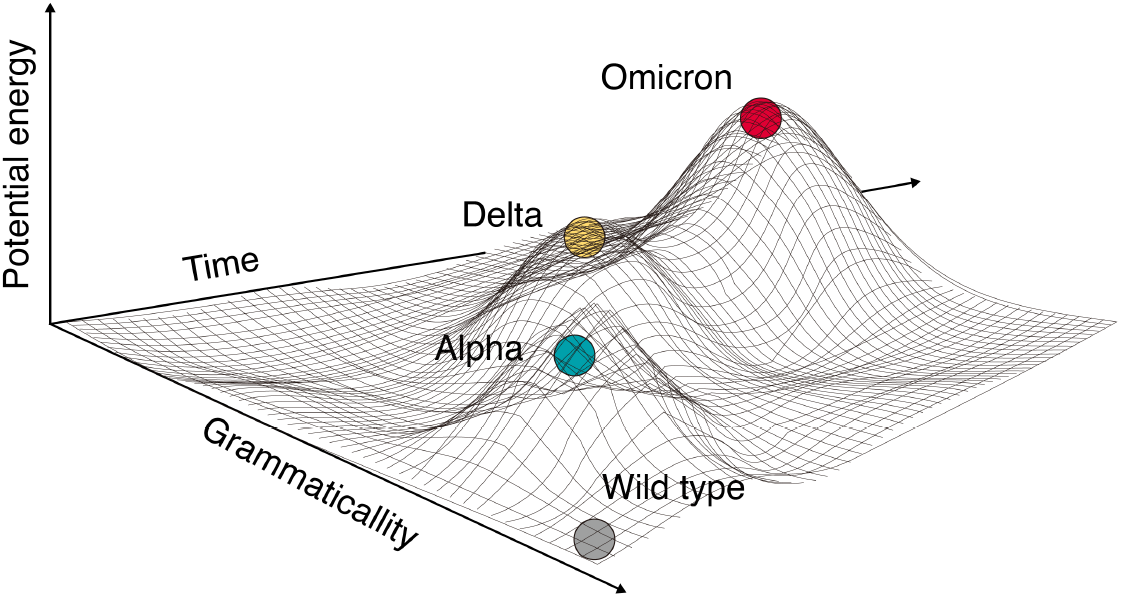
Plot virus genotype fitness landscape. As a two-dimensional hypersurface, the genotype-fitness landscape can plot in a three-dimensional mapping space with “semantics”, “syntax”, and time as axes. Starting from the sampled data, we fitted a quadric surface function and drew a sketch of the Genityp-Fitness landscape, in which only three virus lineages, including Omicron, were shown.

## Extended Data Tables

**Extended Data Table 1.**
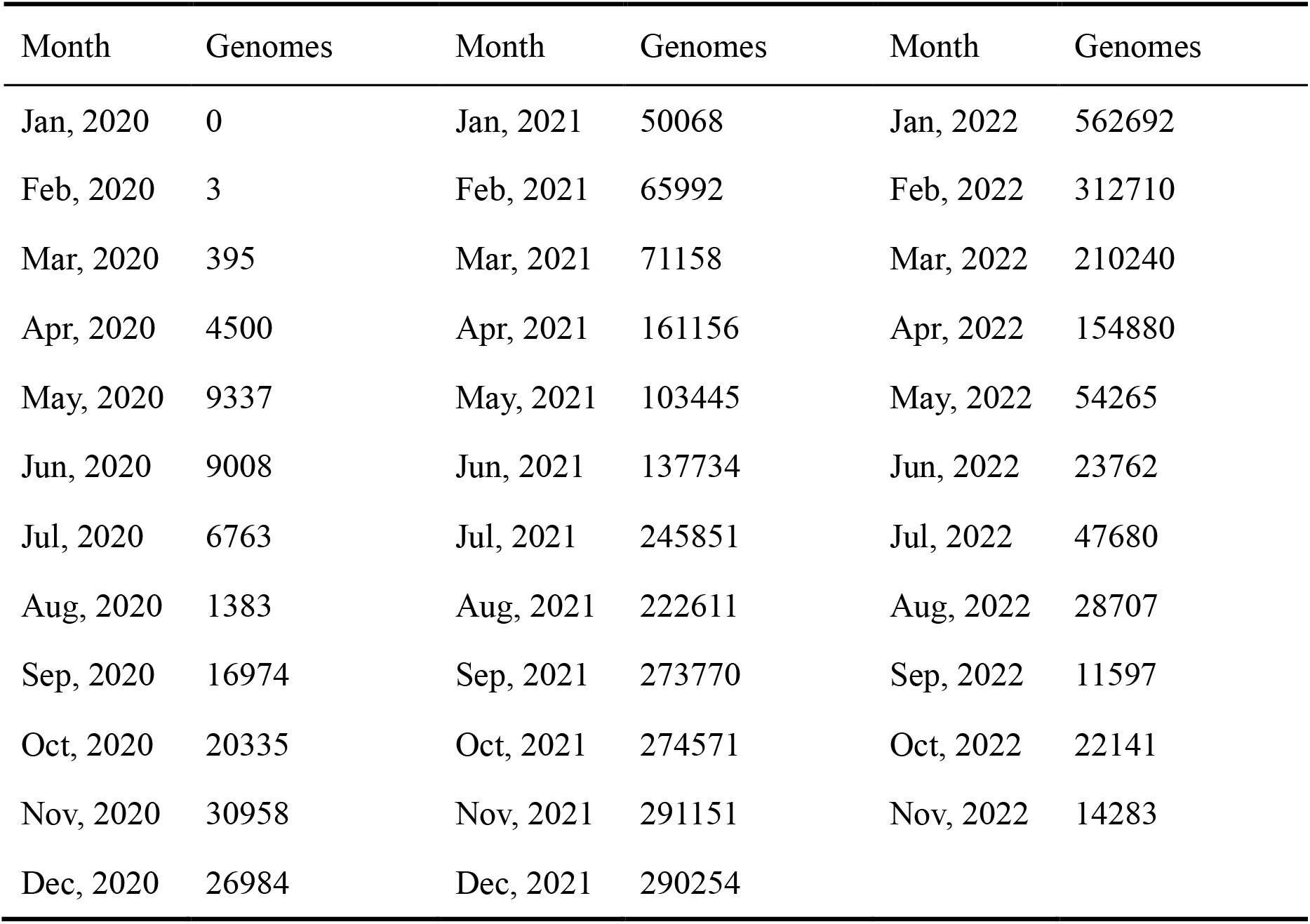
Summary of SARS-CoV-2 Spike protein sequences from the UK involved in this study. These data are used by the model to predict “immune escape” mutation and to calculate the *R*_*0*_ of the virus lineage.

| Month | Genomes | Month | Genomes | Month | Genomes |
| --- | --- | --- | --- | --- | --- |
| Jan, 2020 | 0 | Jan, 2021 | 50068 | Jan, 2022 | 562692 |
| Feb, 2020 | 3 | Feb, 2021 | 65992 | Feb, 2022 | 312710 |
| Mar, 2020 | 395 | Mar, 2021 | 71158 | Mar, 2022 | 210240 |
| Apr, 2020 | 4500 | Apr, 2021 | 161156 | Apr, 2022 | 154880 |
| May, 2020 | 9337 | May, 2021 | 103445 | May, 2022 | 54265 |
| Jun, 2020 | 9008 | Jun, 2021 | 137734 | Jun, 2022 | 23762 |
| Jul, 2020 | 6763 | Jul, 2021 | 245851 | Jul, 2022 | 47680 |
| Aug, 2020 | 1383 | Aug, 2021 | 222611 | Aug, 2022 | 28707 |
| Sep, 2020 | 16974 | Sep, 2021 | 273770 | Sep, 2022 | 11597 |
| Oct, 2020 | 20335 | Oct, 2021 | 274571 | Oct, 2022 | 22141 |
| Nov, 2020 | 30958 | Nov, 2021 | 291151 | Nov, 2022 | 14283 |
| Dec, 2020 | 26984 | Dec, 2021 | 290254 |  |  |

**Extended Data Table 2.** Indels predicted with the sequences from August 2021 and September 2021 (before the Omicron emerging) With SARS-CoV-2 Spike protein sequences of September and October of 2021, a total of 7 mutations (referring to Wuhan-Hu-1) were predicted with the model. These mutations were subsequently identified in Omicron variants.

| Indel | Classification | Omicron variants |
| --- | --- | --- |
| H69del | Confirmed<br>in<br>Omicron variants | BA.1, BA.4, BA.5, BQ.1, B.1.525 |
| V70del |  | BA.1, BA.4, BA.5, BQ.1, B.1.525 |
| G142del |  | BA.1, BA.2, BA.4, BA.5, BA.2.12.1, BA.2.75, BQ.1, XBB |
| V143del |  | BA.1 |
| Y144del |  | BA.1, XBB, B.1.525 |
| N211del |  | BA.1 |
| H66del | Novel | NA |
| A67del (next to H69del) |  | NA |
| I68del (next to H69del) |  | NA |
| Y145del (next to G142del) |  | NA |
| L212del |  | NA |
| S477del |  | NA |
| T478del |  | NA |
| A684del |  | NA |
| R685del |  | NA |
| S686del |  | NA |
| V687del |  | NA |
| A688del |  | NA |

**Extended Data Table 3.** Key resources.

| REAGENT or RESOURCE | Source | IDENTIFIER |
| --- | --- | --- |
| Software and algorithms |  |  |
| Python (v3.8) | Rossum et al. [2] | https://www.python.org/ |
| Biopython (v1.76) | Cock et al. [3] | https://biopython.org/ |
| PyTorch (v1.12.1+cu102) | Paszke et al. [4] | https://pytorch.org/ |
| Matplotlib (v3.4.1) | Hunter et. al. [5] | https://matplotlib.org |
| NLTK (v3.6.2) | Klein et al. [6] | https://www.nltk.org/ |
| NumPy (v1.21.1) | Harris et al. [7] | https://numpy.org/ |
| Pandas (v1.1.4) | McKinney et al. [8] | https://pandas.pydata.org/ |
| Scipy (v1.3.1) | Virtanen et. al. [9] | https://scipy.org/ |
| Scikit-learn (v0.21.3) | Pedregosa et al. [10] | https://scikit-learn.org |
| Fairseq (v0.12.2) | Ott et al. [11] | https://github.com/facebookresearch/fairseq |
| Flax (v0.5.2) | Heek et al. [12] | https://github.com/google/flax |
| JAX (v0.3.15) | Bradbury et al. [13] | https://github.com/google/jax |
| Optax (v0.1.3) | Babuschkin et al. [14] | https://github.com/deepmind/optax |
| Transformers (v4.20.1) | Wolf et al. [15] | https://github.com/huggingface/transformers |
| R (v4.2.2) | R Core Team [16] | https://cran.r-project.org |
| R package: tidyverse (v1.3.1) | Wickham et al. [17] | https://www.tidyverse.org/ |

## Supplementary Information

### Supplemental Note 1: The formal expression of fitness in Encoder-Decoder Seq2Seq framework and the P-E theorem

#### 1. Mathematical description of the average and per-lineage fitness of the virus population

From the basic principles and definitions of population genetics [18], it can be known that the mean fitness 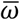 of the virus population (the total number of viruses is *N*) containing *K* virus lineages can be expressed by the following formula:

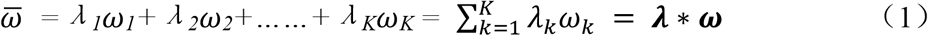

Here, ***λ*** = (*λ*_1_, *λ*_2_, … …, *λ*_*k*_), *λ*_*k*_ is the occurrence frequency of the *k*_*th*_ virus lineage in the whole population, which is an observable measure related to amino acid substitution characteristics, and *ω=*(*ω*_1_, *ω*_2_, … …, *ω*_*k*_), *ω*_*k*_ is the mean fitness of the *k*_*th*_ virus lineage. Apparently, the fitness of the virus population *ω=ω*(***x***) is a very complex high-order function of a random variable ***x*** that describes the variation state of a virus genome sequence sites, and ***x =*** (*x*_*1*_, *x*_*2*_, …, *x*_*i*_, …, *x*_*L*_) represent *L* variation sites. Thus, each *ω*_*k*_ can involve the epistatic interaction of multiple mutation sites in the input sequence [19] as following:

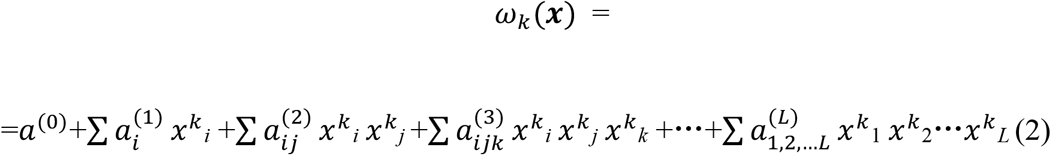

According to the above definitions, *λ*_*k*_*ω*_*k*_ is a component of the fitness of the *k*_*th*_ *>*virus lineage. Given we take the Wuhan lineage as a reference [Wuhan-Hu-1, GenBank: MN908947.3], *ω*_*k*_ can also regard as the relative fitness of the *k*_*th*_ virus lineage. Usually we also call ω_k_ as the relative basic reproduction number (*R*_0_) of the k_th_ virus lineage. The basic definition of the average fitness given by formula (1) is still valid for virus lineages or sub-lineages with a sufficient number of viruses.

On the other hand, we can define a function *P*(***x***) as a state probability distribution of the sequence’s variation states in real space. This variation causes changes in viral fitness. The *P*(***x***) can have an expanded form using a Gaussian Mixture Model (*GMM*).

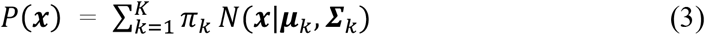

*π*_*k*_ is the mixture coefficient, *N*(***x***|***µ***_*k*_, ***Σ***_*k*_) is the probability density function of a multivariate normal distribution with mean ***µ***_*k*_ and covariance matrix ***Σ***_*k*_. These coefficients *π*_*k*_, ***µ***_*k*_, and ***Σ***_*k*_ can obtain indirectly through the *EM* algorithm or deep neural network.

Now let’s express the formula (1) in terms of the mathematical expectation of a probability density distribution function. According to formulas (3), the formal representation of the average fitness of the virus population can be obtained as follows:

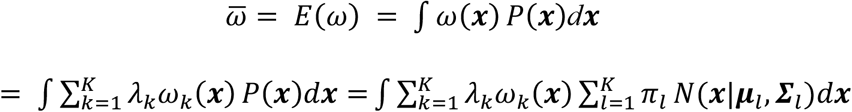

Here, 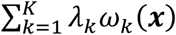 is the linear expansion result of the population fitness *ω*(***x***) according to virus lineage.

Given the independent and identically distributed (*IID*) traits of the virus lineage in the “ideal classification state”, wherein the core of an “ideal classification state” is to attain statistically minimal overlaps among each sub-distribution derived from the expansion of the standard normal distribution function, and the quasi-orthogonal attributes of the base distribution in *GMM* expansion during ideal classification, it is evident that:

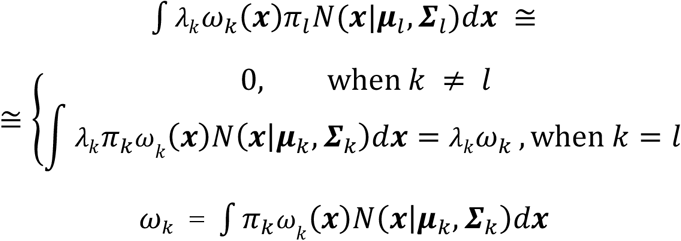

In virus lineage level, *ω*_*k*_ is the fitness of the *k*_*th*_ virus lineage, and in virus population level, *λ*_*k*_= *N*_*k*_*/N, N*_*k*_ is the number of viruses in the *k*_*th*_ virus lineage, *N* is the total number of the virus population, so, 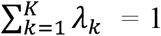, and under this condition conbining with the *EM* algorithm, we get formula (1) again.

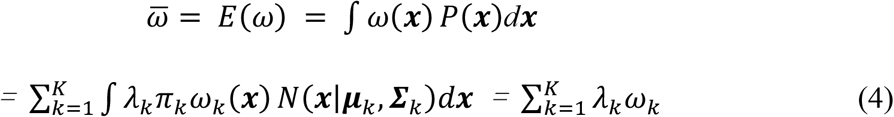

Formula (4) is a representation of the probability distribution of formula (1) in real space. The per-lineage fitness *ω*_*k*_ in real space can be derived when repeating the processes (3) and (4) for the whole individual strain of the *k*_*th*_ virus lineage and in the population leve,

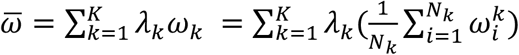

and per-lineage fitness:

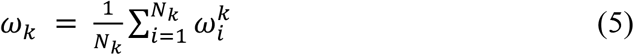

Formula (5) is consistent with the definition by F. Obermeyer et al in their work [20]. According to the basic principles of population genetics [18] that the fitness of a virus lineage can be considered to be determined by the dominant virus cluster in the lineage. Let *ρ*_*k*_ be the occurrence frequency of the dominant cluster in virus lineage when sampling virus strains, and *ρ*_*k*_ = *n*_*k*_/*N*_*k*_, *n*_*k*_ is the total number of samples taken from the dominant cluster in the *k*_*th*_ virus lineage. For the convenience of practical calculation, we can redefine the per-lineage fitness through the contribution of the dominant virus cluster of a lineage as follows:

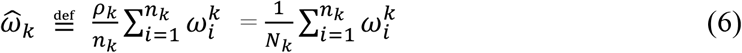

Here, Because of 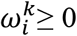, formula (6) can be rewritten as follows for the logical needs of the following proof:

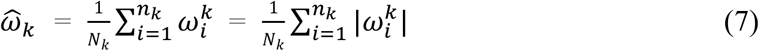

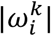 represents the contribution of the sampled virus strain to the fitness of the *k*_*th*_ virus lineage. Now the problem of finding the per-lineage fitness is transformed into finding the dominant cluster in virus lineage and calculating the variable 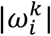.

#### 2. Fitness calculation based on the Bayes’ theorem and deep learning model of the Encoder-Decoder Seq2Seq architecture

According to the definition of the virus fitness (1) and formula (5),

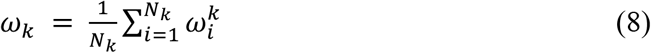

is the per-lineage fitness of the virus population and a mathematical expectation of the contribution of a single virus strain to the fitness. Referring to formula (3), (4) and (5), and let ***λ*** = (*λ*_1_, *λ*_2_, … …, *λ*_*k*_) and ***ω***(***x***) = (*ω*_*1*_, *ω*_*2*_, ……, *ω*_*k*_) are vectors, then

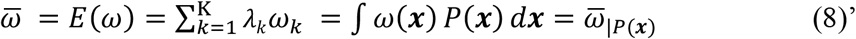

The subscript *P*(·) represents the state probability distribution corresponding to virus population. As long as we find a suitable functional form of *ω*(***x***) and a corresponding probability distribution *P*(***x***), we can obtain the fitness of viruses by calculating their mathematical expectations by the formula (8), then get the average fitness of the virus population and per-lineage fitness of virus lineages by formula (4) and (6). We can now try to do this within the context of deep learning, despite the fact that it is typically difficult to obtain and calculate ω(***x***) and *P*(***x***) directly from real-world data with noise.

Let *P* (***z***|***x***) and *P*(***z***) be the hidden state probability distribution in encoder-decoder spaces, and *P*(***x***) and *P*(***x***|***z***) be the prior and posterior probability related to the input and output of the model. The relationship among *P*(***x***), *P*(***z***|***x***), *P*(***x***|***z***) and *P*(***z***) is given by Bayes’ theorem:

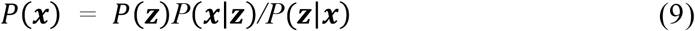

The function relationship defined by formula (1) still holds when *P*(***x***) maps to *P*(***z***) *P* (***x***|***z***) */P* (***z***|***x***). When the prior probability *P*(***x***) is unknown, obtaining the fitness of virus lineages in the encoder or decoder space is transformed into finding the hidden state probability distributions *P*(***z***|***x***), *P*(***z***), *P*(***x***|***z***) and the counterpart 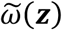 of *ω*(***x***).

Under the Encoder-Decoder Seq2Seq framework together with Bayes’ theorem and Universality Theorem [21], because

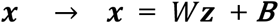

we have

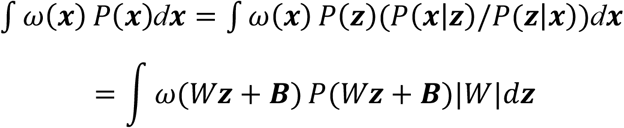

Here,

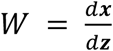

is the Jacobian matrix, and 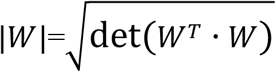 is a determinant of Jacobian matrix that is the sum of combinations of products of elements in different positions of all the columns of matrix *W*. and through the encode-decode architecture’s deep learning process, coupled with the Seq2Seq property, we can derive a parameterized representation of Bayes’ theorem:

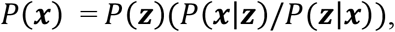

At this juncture, considering the Universality Theorem, convergence prerequisites of deep neural networks and the linear transformation characteristics of ***x****=Wz+B* in the output layer and the neighboring hidden layer, the equation

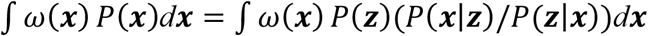

holds. At this juncture, employing the linear transformation ***x****=W****z****+B*, the integral of x is translated into the integral of ***z***, yielding the equation

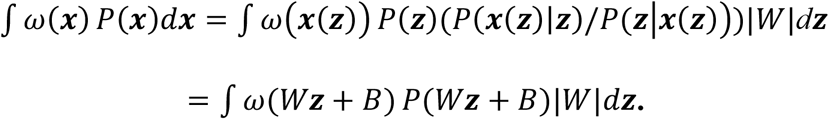

if we set:

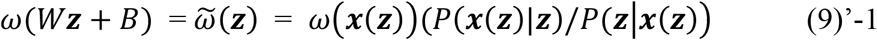

This equation can be rewritten as:

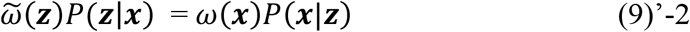

Equation (9)’-2 implies that if 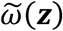 is a topological mapping of *ω*(***x***) in the hidden space within deep neural networks, then the product of the function *ω*(***x***) of the microstate variable ***x*** and the conditional probability distribution of ***x*** in the real space equals the product of the function 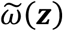 of the latent variable ***z*** and the conditional probability distribution of ***z*** in the hidden space. If we take into account the topological properties of encode-decode Seq-Seq deep neural networks, it is evident that this equation holds true. It follows naturally.

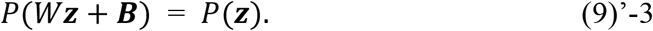

Consequently, we get finally

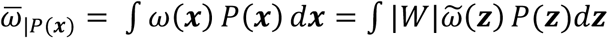

Given the quasi-orthogonal property inherent in the base distribution of Gaussian Mixture Model (*GMM*) expansion for optimal classification, let *β*_*k*_ represent the mapping of |*W*|_*k*_ for the *k*_*th*_ virus lineage, and 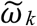 denote a hidden space state function associated with the fitness of the *k*_*th*_ virus lineage. Forming vectors ***β*** = (*β*_1_, *β*_2_, … …, *β*_*k*_) and 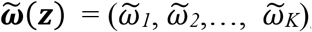, the relationship

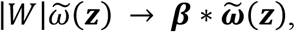

holds true. Then we get

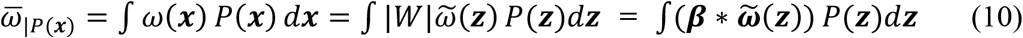

When a virus lineage contains enough virus strains, let *ω*(***x***) = *ω*_*k*_(***x***), *P*(***x***) = *P*_*k*_(***x***), and combining the formula (4), the following formula still holds,

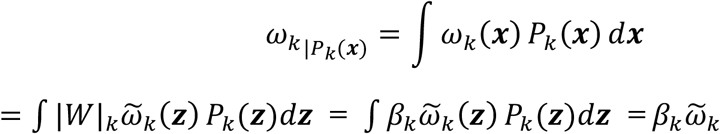

From formula (10) and see the logic for deriving formula (4), and combining formula (5), we have

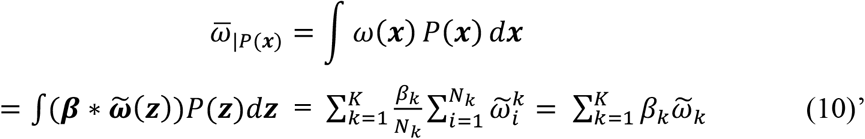

Formula (10)’ shares an identical structure with formulas (8) and (8)’. Hence, it is reasonable to equate ***β*** = (*β*_1_, *β*_2_, … …, *β*_*K*_) = ***λ*** = (*λ*_1_, *λ*_2_, … …, *λ*_*K*_),. Consequently,

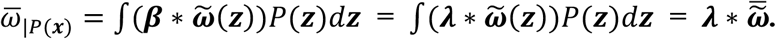

This leads to the final expression:

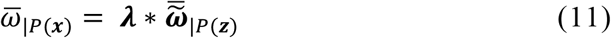

Formula (11) is a representation of the probability distribution of formula (1) in an embedded space. Because of the hierarchical structure of the population, lineage and cluster of the virus, referring to the formula(1), (3), (4), (6) and (10) and based on the basic principles of population genetics [18], the elements of the vector ***λ*** are the corresponding elements of mixture coefficient in formula (3). According to the formula (6) the element *λ* also associated with the occurrence frequency of a dominant cluster in virus lineage when sampling virus strains in a virus lineage and is a macroscopic parameter that links the hidden space with the actual space.

Under Encoder-Decoder Seq2Seq framework, *P*(***z***) is a prior probability distribution of hidden space. Now our task is to find a 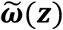 corresponding to the *ω*(***x***) in hidden space.

Based on formula (10), we know that *ω*(***x***) and 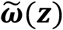 belong to two different spaces. One is the real-world space, where the input is the mutation information of the virus sequence ***x***. The *ω*(***x***) is a very complex high-order function involving the epistatic interaction of multiple mutation sites in the input sequence (see formula (2)) However, based on the embedded mode of deep learning, the learning mechanism of fully connected network linear transformation and dimension reduction characteristics, the 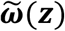 in hidden space can be a low-order simple quasi-linear function of a latent variable ***z*** to achieve accurate fitness calculation. Therefore, the formula (10) and (11) show that the fitness of the virus lineage can be computed as a mathematical expectation according to a hidden state probability distribution under the Encoder-Decoder Seq2Seq framework if we can find a reasonable embedded representation of the related microscopic genome sequence variation.

In this way, to calculate the fitness of the virus population, according to formulas (10) and (11), we need to establish a Seq2Seq deep learning model under the Encoder-Decoder Seq2Seq framework to realize our calculation of the fitness of the virus population.

Considering that the fitness of the virus population is an observable macroscopic phenotype, and genome sequence variation is related to microscopic genotype change, the meaning of equation (11) can be generalized, and the following phenotypic embedding (P-E) theorem in the biological field can be proposed:

#### Inference

The P-E theorem: “An observable macro-biological phenotype can be computed under the Encoder-Decoder Seq2Seq framework if we can find a reasonable embedded representation of the related microscopic genotype.”

### Supplemental Note 2: The fulcrum of building model system and the introduction of CSC scores

Our goal is to find virus mutations that cause a high degree of semantic change but are grammatically acceptable, from which to identify and predict “immune escape” mutations, and then calculate the fitness of virus lineages. Because of B. Hie’s pioneering work [4], and the need for a clear and similar pivot to construct our theoretical system, in addition to some modifications required by our model, we adopted the basic formulation of DNA base or amino acid embedding given in [22], and it is paraphrased as follows:

For a virus sequence defined as 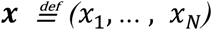 such that *x*_*i*_ ∈ 𝒳*i* ∈ [*N*], where 𝒳 is a finite set of characters, ***x*** ∈ *V, V* is virus population set and ***x***_*k*_ *∈ V*_*k*_, *V*_*k*_ is virus lineage set. Let 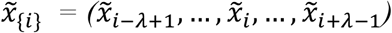 denote mutations around position i and the mutated sequence is 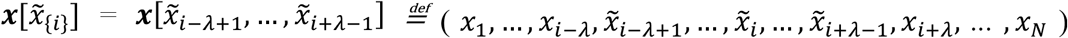, here, *1<=λ<=*Length(protein sequence) *\** 1*5%* and for SARS-CoV-2 spike protein 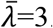. For brevity, we will use 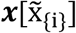 instead of 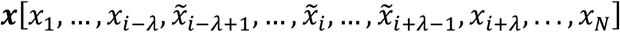 in the following. The syntax for a mutated sequence ***x*** is described as a probability distribution function:

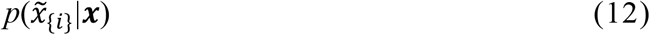

Then, 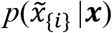 is close to zero if 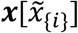 does not conform to the grammar rules, and 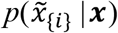 is close to one if 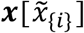 conforms to the grammar rules. In a language composed of a given sequence, violation of these grammatical rules results in a loss of grammaticality of the sequence. To establish the correlation between the above definition and the natural language model, a latent variable distribution function 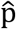 and the mapping 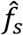 can be introduced, and for all *i*∈[*N*], let the latent variable 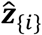 is encoded from the context 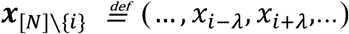, in which the viral genome mutates at position (*x*_*i* − *λ*_, *…, x*_*i* **+** *λ*_). In our model CoT2G-F, 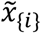 in ***x***_*[N]\*{*i*}_ is randomly masked, and then in the original position 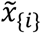 re-gives the character according to the “grammar rules” learned by the model, besides, the multiple consecutive sites (*λ>* 1) have been masked and they are treated as a “span” and a single unique mask token is used to replace the entire span. Thus, we have

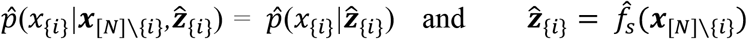

We define:

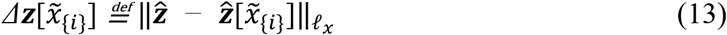

and 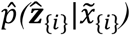 as the mapping representation of 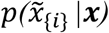 into the encoding hidden state space under our model, and 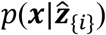 is the mapping representation of 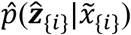 in the decoding recovery state space and ***x*** is the output sequence of our Se2Seq model,

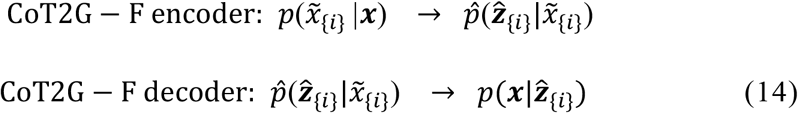

Here [22] defines

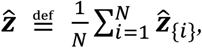

which is the average of semantic embeddings at all positions. Formula (2) shows that 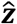 is a reference value for embedded “semantic” changes, and what virus lineage is chosen as a reference has biological significance. In our work, the Wuhan lineage was selected as a reference. 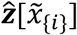 is the sequence embedding that does not contain the variant site 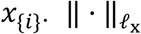 is the *𝓁*_*x*_ norm, x = 1, 2, and 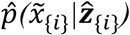 can be obtained through model training.

In our model CoT2G-F (see supplemental note 2)

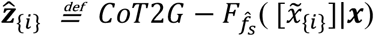

When there are multiple mutations in the sequence, as defined

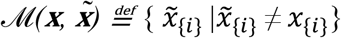

According to formula (3), the hidden state probability distribution function of the decoding space is rewritten as

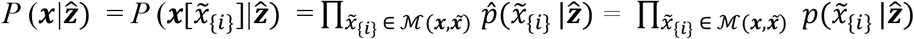

In our CoT2G-F model, we can further get

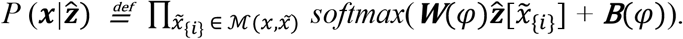

Where ***W*** and *B* are both the parameters the model needs to learn and 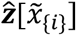 obey a hidden state probability distribution. In this way, when judging semantic and grammatical changes and identifying and predicting “immune escape” mutational patterns, the “semantic” change 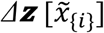 remains unchanged, and the rules from [22] still hold:

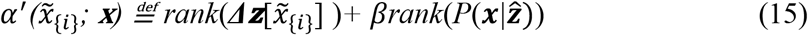

The distance 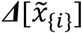 in the embedding space describes the semantic change, and the hidden state probability distribution 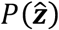 and the related posterior probability 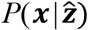 gives the grammatical rules that constrain the sequence composition to be grammatical.

Based on the principles of evolutionary biology, fitness is a concept of change. If the genome does not change, there is no fitness change. Therefore, the “grammaticality” of the genome composition will not directly affect the fitness of the virus. If the genome composition is already following “grammaticality”, only the “semantic” changes corresponding to the sequence variations affecting the genome function contribute to the fitness of the virus. Referring to the study of [22], we think that the first term 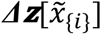 on the right side of equation (4) is a variable related to the fitness of the virus under the condition that its sequence composition is grammatical.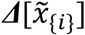 can also regard as a conditional “semantic” change 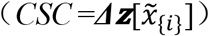 score. Late we will show that the per-lineage fitness can derive from the measure *CSC* with a hidden state distribution 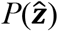 and 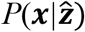 in the decoder space (see supplemental notes 3 and 4).

### Supplemental Note 3: Detailed description of the CoT2G-F model

For our CoT2G-F model, we use a vanilla encoder-decoder Transformer [23], which has emerged as a powerful general-purpose language model architecture for representation learning and generative modelling, outperforming recurrent and convolutional architectures in natural language settings.

We take as input amino acid character sequences. Protein sequences are composed of amino acid characters and natural language sequences are composed of words that can both be processed in language models based on different vocabulary. Researchers [22,24,25] have demonstrated that protein language models trained in diverse sequences can learn the structure and function of proteins, the metric structure in the protein representation space that accords with organizing principles at scales from physicochemical to remote homology.

Language mode is commonly trained with two steps: pre-train and fine-tune, which corresponds to neutral evolution and environmental selection during virus evolution. The first stage is pre-train, where a high-capacity language model is trained on a large amount of unlabeled data, and such a language model can capture general semantic and syntactic information from sequence data alone. Then for the second fine-tuning stage, models apply the first stage pre-trained language model to a particular task, which means a task-specific input and objective are introduced. In other words, the model is fine-tuned on the task data from pre-trained parameters. In our model’s pre-training step, we not only leverage the 20 million GISAID database of virus sequences but also introduce a co-attention mechanism to capture the spatial correlations of virus evolution. The self-attention mechanism in the vanilla Transformer only focuses on the internal context information of the sequence itself. The query, key, and value in the attention function are all from the sequence itself. By contrast, in our CoT2G-F model, the co-attention mechanism allows our model to introduce Spatio-temporal neighbour sequences that participate in the key and value of the attention function. Then the learned representation from pre-training has a multi-scale organization reflecting structure from the level of biochemical properties of amino acids to the remote homology of proteins. A spatial correlation reflecting the sequence structure composition rules of virus genomes. Both properties contribute to our downstream tasks: identify the immune evasion sites of viruses and calculate their relative fitness only from sequence information.

Pre-training is done using a denoising objective, where some parts of the amino acid characters are randomly replaced with a special token, and the model needs to predict the identity of those amino acid characters, which is also called the “masked language modelling” objective. Inspired by BERT’s [26] “masked language modelling” objective and the “word dropout” regularization technique [27], we design an objective that randomly samples and drops out 15% of consecutive amino acid characters in the input protein sequence, during the pre-train step, the multiple consecutive amino acid characters have been masked and they are treated as a “span” and a single unique mask token is used to replace the entire span. All spans of discarded amino acid characters are replaced by a single marker, each assigned a mask token ID that is unique to the sequence of amino acid characters. These mask token IDs are special tokens which are added to our new vocabulary, we choose to mask contiguous spans of amino acid characters and only predict discarded spans that can reduce the computational cost of pre-training. The loss function to predict the missing amino acid characters from the corrupted sequence is followed:

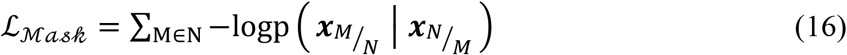

For each sequence *x*, sample a set of indices N contiguous spans of amino acid characters to mask, replacing the true amino acid characters with the mask token IDs. We minimize the negative log-likelihood of the unmasked amino acid characters x_M/N_ given the masked sequence x_N/M_ as context for every masked token ID.

To discover the implicit alignments between two spatial contiguous sequences of protein sequences, we introduce a co-attention mechanism. For each sequence ***x***_(*‘,^*)_ at t month slice,***x***_(*t,p* −1)_ and ***x***_(*t,p* **+**1)_ is the spatial pre-order and follow-up contiguous sequences of ***x***_(***t***,***p***)_, [:] means concatenate two sequences, so the

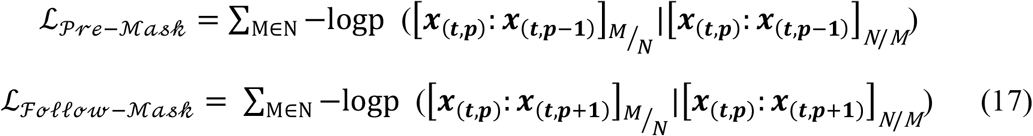

The pre-training is done using a combination of pre-order denoising objective and follow-up denoising objective, to speed up the convergence of the model, we adopted a staged training strategy, i.e. our method first set *α*_*𝒫𝓇ℯ*−*ℳ𝒶𝓈𝒽*_ = 1, *α*_*ℱℴ𝓁𝓁ℴ𝓌*−*ℳ𝒶𝓈𝒽*_ = 0, then set *α*_*𝒫𝓇ℯ* − *ℳ𝒶𝓈𝒽*_ = 0, *α*_*ℱℴ𝓁𝓁ℴ𝓌* −*ℳ𝒶𝓈𝒽*_ = 1 and continue training on the checkpoint of the previous model.

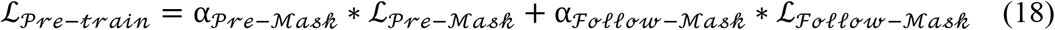

In the fine-tuning step, we follow the Constrained Semantic Change Search (CSCS) criterion proposed by B. Hi [22], CSCS is a sorting algorithm that sorts the semantic and grammatical changes of a base sequence to multiple sequences that may be mutated and find the mutation sequences that are most prone to ‘immune escape’ from highest to lowest. We denote such a base sequence as ***x*** and the mutated sequences as 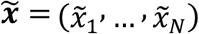, we assume there are *N* mutant sequences, for one mutant sequence 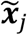, which consists of amino acid characters in 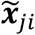 that disagree with those at the same position in the base sequence ***x***, which we denote as

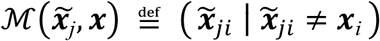

All possible mutations 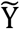 are given priority based on the corresponding values of CSCS, from highest to lowest.

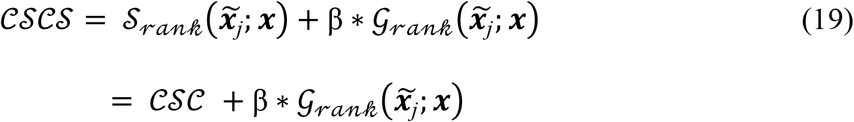

where 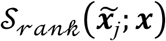 and 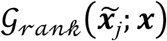 denote semantic and grammatical rank respectively, the consistently well-calibrated around *β* =1 equally weighting both semantic and grammatical change), which we used in our experiments.

The semantic change is measured by the following:

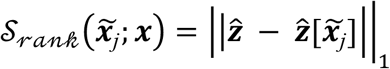

Where 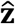 is the hidden states of the Transformer’s last decoder block and || ⋅ ||_1_ is the *𝓁*_1_ norm.

The distance in the embedding space encodes semantic changes, while probabilistic emanations encode grammatical changes:

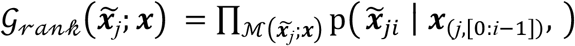

where *p* is the mutation probability that the *i* site corresponds to the 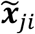 is mutated to the amino acid corresponding to the *i* site in ***x***, and the grammatical change is equal to the product of the probabilities of the individual mutations.

Following the above *CSCS* criterion, we perform protein sequences set slicing monthly, select the virus strain with the largest semantic change but conform to the grammar in the next month, these selected protein sequences can “immune escape” most likely, from which the labelled sequence-to-sequence dataset during fine-tune step is constructed. The objective of fine-tuning step is as follows:

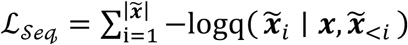

For each sequence ***x*** at one month slice, the model needs to generate a list of target sequences, donated as 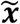, which is the virus strain that may emerge “immune escape” mutations in the next month, the 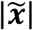 means all sites of 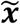, and due to the autoregressive property of the decoder, when predicting 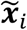, the sites before the i site can also be introduced, i.e.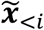. We need to independently minimize the negative log-likelihood of the amino acid 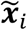 given the ***x*** and 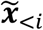 as context. Then based on pre-trained language model parameters, iterative training and dynamic inference of immune evasion strains are performed simultaneously, thus, a temporal correlation reflecting the selection mechanism is introduced at this fine-tuning step.

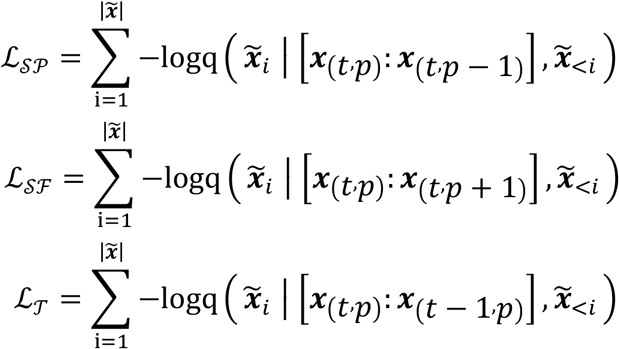

We endow the language model with the ability to predict new immune escape sites arising from environmental selection and time migration following a neutral evolution event. Fine-tune is done using a combination of three objectives, refer to the staged training strategy mentioned above, we set *α*_*𝒮𝒫*_ = 1, *α*_*𝒮ℱ*_ = 1 and *α*_*𝒯*_ = 1 successively.

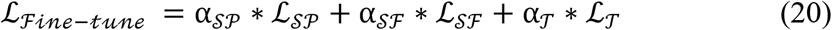

### Supplemental Note 4: The formal logic of the Genotype-Fitness landscapes and *R*_*0*_ calculation

#### 1. The “immune escape” ability score *CSC*

B. Hie et al. proposed a semantic-grammatical criterion *CSCS* to determine the ability of the virus “immune escape”, which is expressed by the following formula [22]:

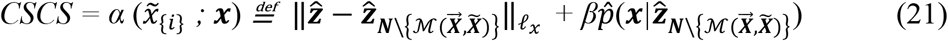

From the logic of evolutionary biology, combined with our model,

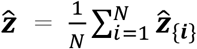

is the mean of all possible “semantic” changes in a sequence of *N* bases. If we select the Wuhan lineage as the reference [Wuhan-Hu-1, GenBank: MN908947.3], 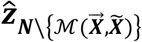 is an absolute semantic change of a virus strain, and 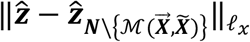 is a relative semantic change of a virus strain according to the Wuhan virus lineage. In this way, a conditional semantic change score (*CSC*) under the condition that the sequence composition conforms to the grammar can be defined as:

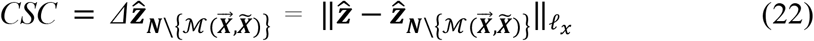

and we can rewrite (21) as follows

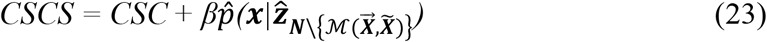

*CSC* score is the measurement of the “immune escape” ability of the virus strain referring to the Wuhan lineage when the virus mutates. Formula (1) gives the fundamental definition of the mean fitness of the virus population and the fitness of the virus lineage. It should note that this is the fundamental starting point of the definition of fitness in evolutionary biology. Here, starting from the basic definition of fitness and homology in population genetics and considering the computational complexity, we adopt the *𝓁*_1_ norm to define the *CSC* score. Because our model has better avoiding overfitting characteristics (Figure 3).

#### 2. The mathematical expectation of the *CSC* score according to the prior hidden state probability distribution *P*(***z***) is consistent with the formal expression of the average and the per-lineage fitness of the virus population

Based onur proposed Encoder-Decoder Seq2Seq architecture model, the encoder module created an embedded state distribution *P*(***z***|***x***) related to the virus fitness from the input genome sequence ***x***. The decoder module can characterize the “immune escape” of the virus and decode a hidden state probability distribution *P*(***z***) of a latent variable ***z***. *P*(***z***) can also be formally expanded by the Gaussian Mixture Model (*GMM*) kernel function:

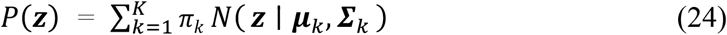

Where *π*_*k*_*N* (***z*** / ***µ***_*k*_, ***Σ***_*k*_) is called the *k*_*th*_ component in the mixture model, corresponding to the probability distribution of the *k*_*th*_ virus lineage, and *π*_*k*_ is the mixture coefficient satisfying *π*_*k*_ ≥ 0, and 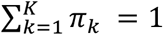.

Definition

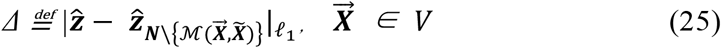

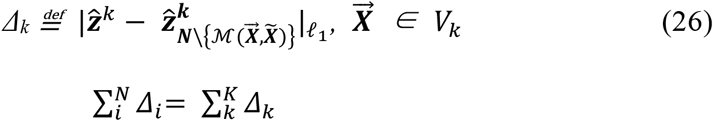

*N* and *K* are the number of virus strains and the number of virus lineages in a virus population respectively. 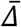 and 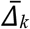 are the expected values of *Δ* and *Δ*_*k*_ according to the whole distribution of viruses and the sub-distribution of the *k*_*th*_ virus lineage in the decoder space, respectively. In our Encoder-Decoder Seq2Seq architecture model, combining formulas (22), (25) and (26), and the quasi-orthogonal characteristics of the base distribution of *GMM* expansion in the ideal classification, the average “immune escape” ability of the virus population can express as:

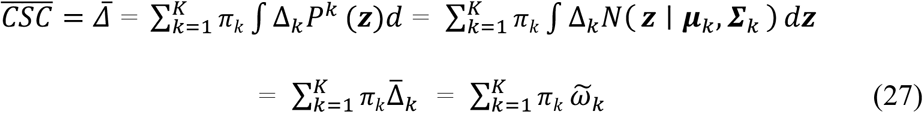

Here, the component *P*^*K*^(***z***) is the prior probability distribution of the *k*_*th*_ virus lineage, and together with the formula (22), the average “immune escape” ability of the *k*_*th*_ virus lineage is:

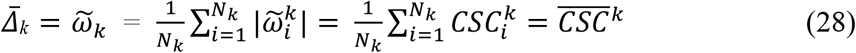

The 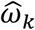 has a mathematical representation consistent with (7). Here,

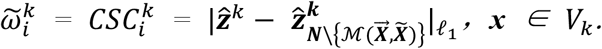

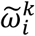 is the “immune escape” ability determined by “semantic changes” of a mutated virus strain in the *k*_*th*_ virus lineage compared with the Wuhan virus lineage, and

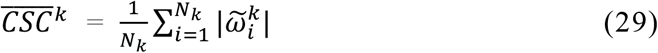

is an absolute mean of the 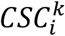, a mean of the cumulative effects of the “immune escape” ability of a virus strain for the *k*_*th*_ virus lineage. We define 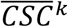 as the average immune escape ability of the *k*_*th*_ virus lineage.

Let us look at the distribution of the absolute mean “semantic” changes computed by sampling different virus lineages (Figure 3B). The trend of this distribution is highly consistent with the fold increase in relative fitness over the Wuhan (A) lineage given by F. Obermeyer, et al. [20]. Let us now consider the following four facts: (i) The model determined that latent variable ***z*** reflected the semantic and grammatical changes of virus sequence site variation and the correlation between variation sites given by formula (2) and [22]; (ii) After the selection of a reference virus lineage (in this case, the Wuhan lineage as the reference lineage), from formula (22) the *CSC* score reflects the degree of semantic change of a virus strain after a new sequence variation, indicating its possible contribution to the fitness related to the immune escape ability of the lineage; (iii) The *CSC* score correctly reflected the changes in immune escape ability caused by virus sequence mutation under syntactic fitness constraints (*CSCS* constraints [22]) (Figure 2B); (iv) The mean value of the *CSC* score as the convolution of “semantic” and “grammatical” terms of *CSCS* criterion correctly reflected the whole immune escape ability of the virus lineage (Figure 3), and the variation trend of the virus across lineages given by mean *CSC* score was consistent with the *R*_*0*_ calculation by the work [20] and general epidemiological observation.

Based on the above four facts, we believe that 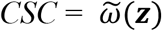 is the corresponding function of *ω*(***x***) in hidden space. It’s a lower-order function of the latent variable ***z***. Based on Bayes’ theorem (9) and together with formulas (1), (6), (22) and (29), and set 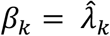, we have

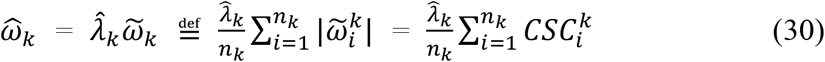

Here, 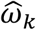 is the per-lineage fitness of the virus and it can be calculated directly under the framework of deep learning, 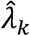 is the occurrence frequency of the dominant cluster in the *k*_*th*_ virus lineage. Thus, formula (27) is the fitness of the virus population. In this way, we prove that the item of the degree of “semantic” change in the *CSCS* criterion proposed by B. Hie, et. al., [22] is related to the per-cluster fitness of the virus given by F. Obermeyer, et al. [20]. Although all the parameters involved in the calculation of the per-lineage fitness 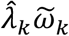 can be computed directly by the model, it is clear that the parameter 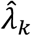 and latent variable ***z*** indirectly reflect the effects of selection. Formula (30) is the concrete realization of the P-E theorem (see formula (11)) under the CoT2G-F model.

We can carefully compare formulas (15), (22) and (30) to find there exists an intrinsic correlation between the semantic change term and the grammatical term. This association can express as a convolution of the semantic variation term with the hidden state probability distribution because the grammar term in our model is the posterior distribution of the hidden state probability distribution. This convolution can also express as a mathematical expectation of the *CSC* according to the hidden state probability distribution *P*(***z***). Interpreted in the sense of convolution, it is precisely the mean value of the accumulation of “semantic” changes of the virus lineage under the constraints of “grammar”. And this mean is the same in form as the fitness calculation formula (15) we got from evolutionary biology, and the calculation formula of the immune escape ability of the *K*_*th*_ virus lineage is deduced.

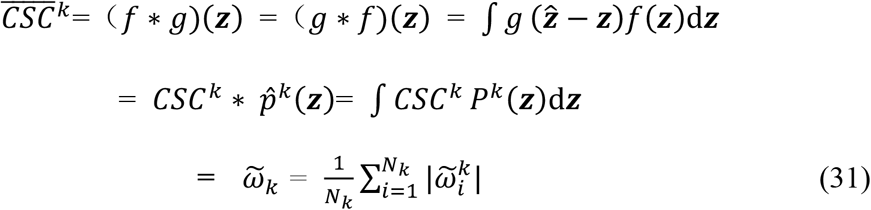

Here,

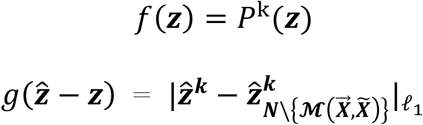

At this point, all we need to do is start from actual data and calculate the correctness of our mathematical framework. Therefore, we need to calculate the per-lineage fitness of the *k*_*th*_ virus in the unit area *R* and unit time *t*_*bin*_ according to the following formula by sampling:

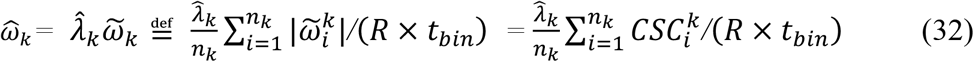

*R* is an area, such as the United Kingdom, and *t*_*bin*_ is a passage time from occurrence to an epidemic of a virus lineage. *n*_*k*_ is the sampling number of the dominant cluster in the *k*_*th*_ virus lineage within the range of (*R × t*_*bin*_), 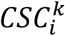 is the conditional semantic change score of the *i*_*th*_ virus strain of the *k*_*th*_ virus lineage, and 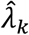 is the occurrence frequency of the dominant cluster in the *k*_*i*_ virus lineage. A reasonable sampling time interval *t*_*bin*_ can be selected for calculating the 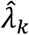 (see supplementary materials, Extended Data Supplementary Figure 3A and 3B). At this time, the problem of finding *R*_*0k*_ of the *k*_th_ virus lineage transforms into computing the mathematical expectation 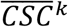 of the 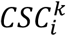 of the virus sequence using the deep learning method according to the following formula:

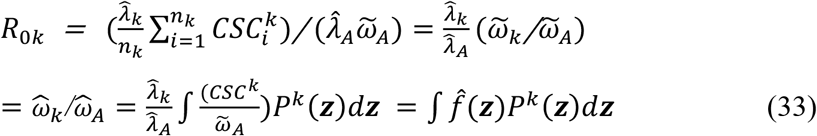

Here,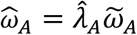 is the fitness of the Wuhan lineage, 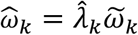 is the fitness of the *k*_*th*_ virus lineage, and the relative basic reproduction number *R*_*0k*_ is the fold increase of the per-lineage fitness 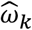 over 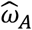. At this point, we finally used the deep learning model to realize the calculation of formula (1) for SARS-CoV-2 virus population. Now the relative fitness or the relative basic reproduction number (*R*_0_) of the virus lineage are precisely computable, which is proportional to the occurrence frequency of the dominant cluster in the virus lineage times the absolute mean of the CSC score according to the hidden state probability distribution.

#### 3. The prediction of immune escape mutation sites and calculation of *R*_*0*_ based on the CoT2G-F model

The methods are significantly different when using the CoT2G-F model to predict immune escape mutation sites and calculate *R*_0_. The detailed steps are as follows:

##### Prediction of immune escape mutations

When predicting immune escape mutation sites, we first need to continuously and randomly mask the amino acids in the input sequence, then based on the *CSCS* criterion proposed by B. Hie (4), we have designated the model output as the protein sequences most prone to immune escape in the training phase. So in the prediction phase, the model would naturally compound this paradigm and output the most likely protein sequences for immune escape in the next time slice. Afterwards, based on the comparison results with the Wuhan sequence, we can determine the possible immune escape mutation sites in the generated protein sequence i.e., we regard the CoT2G-F model as a generation model. After generating various sequence mutations, we use the probability of mutations learned by the model to determine immune escape mutation sites through *CSCS* scoring.

##### *R*_*0*_ calculation

When calculating *R*_0_, we no longer regard the CoT2G-F model as a generation model but as a scoring model. For the input SARS-CoV-2 spike protein sequence, the model does not generate any mutation sites, and only uses the trained model as a feature extractor to get the corresponding sequence embedding. Through the model, according to the Wuhan lineage, each input sequence is given a *CSC* score calculated by the model. Then we calculate the probability average of the *CSC* to obtain the fitness of each lineage and compare it with the fitness of the Wuhan lineage to obtain the relative fitness, getting relative *R*_0_.

In our study, we propose a simple calibration process to obtain the absolute basic reproduction number 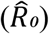 by using a selected observable actual number of the absolute basic reproduction number 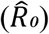 for any lineage of SARS-CoV-2. From formula (33), we can obtain the absolute basic reproduction number 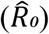 as below

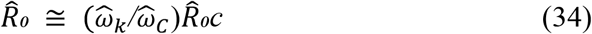

Here, 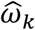 and 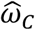 are the relative per-lineage fitness of lineage *k* and *C* according to Wuhan strain, and 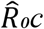 is the actually observed absolute basic reproduction number of the reference lineage *C*. When we select Delta lineage as an observable reference [28,29], 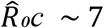. (Figure 4B, Supplemental Note 4, Formula (30)). Our calculated absolute basic reproduction numbers 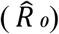 align with the results obtained by F. Obermeyer *et al*. [20] and are also in agreement with the actual and official monitoring results. This result validates our proposed P-E theorem and G-P principle, allowing for the precise computation of the relative fitness and 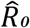 of the virus lineages.

##### Genotype-Fitness land scape derived by the P-E theorem

Based on the P-E theorem and from formula (30), we get

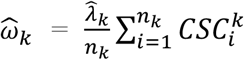

Here, 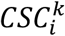 is the conditional semantic change score of the *i*_th_ virus strain of the *k*_*th*_ virus lineage, and is the contribution of a single virus strain to the fitness.

Finally, we can plot the genotype-fitness landscape in the embedded space based on the variation state of the viral genome sequence. Starting from *CSCS* criteria [22], we can regard the *CSC* score as the “immune escape” potential of the virus, which is a function of latent variables and time. As a two-dimensional hypersurface, the genotype-fitness landscape can plot in a three-dimensional mapping space with “semantics”, “syntax”, and time as axes. The peaks on the hypersurface correspond to the genomic variation states that contribute the most to the “immune escape” ability of the virus lineages (see Supplementary Figure 9). We can obtain the fitness of the virus lineage by integrating the surface density of the specific region corresponding to the virus lineage on this hypersurface (Figure 4A).

#### 4. G-P principle: Genotype-phenotype landscapes in embedding space based on the deep learning framework

In equation (33), the *R*_*0k*_ on the left side is a macroscopically observable biological variable, and

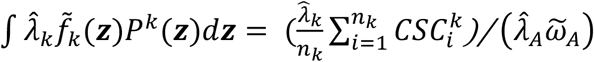

on the right side is a mathematical expectation of a function 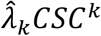 of the latent variable ***z***, which is related to genome sequence variation, according to the hidden state probability distribution *P*(***z***). From equation (8) and (33), we can see that deep learning can help us build a bridge between Genotype-Phenotype, and we can draw a natural inference:

“A macroscopically observable biology variable 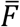 can express as a mathematical expectation of a microscopic state function *f*(***x***) according to a state probability distribution *P***(*x*)**”.

This statement we call the G-P (Genotype-Phenotype) principle, it can represent by the following equation:

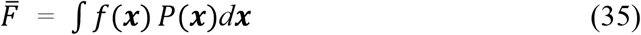

and according to the P-E theorem (11), the formula (35) is equivalent to

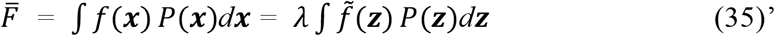

Variable ***z*** is a latent variable, *λ* is a macroscopic parameter that links the hidden space with the actual space. 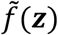 is a function corresponding to *f*(***x***) in hidden space and *P*(***z***) is a prior probability distribution in hidden space, both of 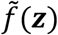 and *P*(***z***) can be calculated by deep learning.

In recent years, many research works have been devoted to discovering the intrinsic theoretical relationship and mathematic basis of deep learning theory and statistical mechanics [24,25]. We know that the core concept of statistical mechanics is: “macroscopic physical observables can characterize as ensemble averages of functions of corresponding microscopic variables”, which can be commonly expressed by the following formula:

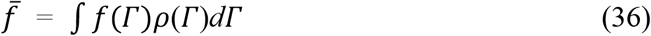

Here: 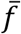 is the macro-observable physical quantity, such as temperature, air pressure, etc., *f*(*Γ*) is the function of the micro-state space variables, and ρ(*Γ*) is the ensemble distribution function of the micro-state space. We see that equations (35) and (36) have the same form. Referring to formulas (10) and (11)), it is clear the G-P principle is an important corollary of the P-E theorem.

Starting from the calculation of virus fitness, based on the basic mathematical framework of deep learning theory, we obtained the genotype-fitness landscape in embedding space. Current work is a practical example that conforms to the G-P principle. It opens up a new field for applying deep learning theory in biological research, giving a new route for the interpretability of models, and presenting theoretical guidance on establishing a genotype-phenotype landscape.

